# Synaptic mitochondria are critical for hair-cell synapse formation and function

**DOI:** 10.1101/671701

**Authors:** Hiu-tung C. Wong, Qiuxiang Zhang, Alisha J. Beirl, Ronald S. Petralia, Ya-Xian Wang, Katie S. Kindt

**Author notes:** **Corresponding author** Katie Kindt.

## Abstract

Sensory hair cells in the ear utilize specialized ribbon synapses. These synapses are defined by electron-dense presynaptic structures called ribbons, composed primarily of the structural protein Ribeye. Previous work has shown that voltage-gated influx of Ca^2+^ through Ca_V_1.3 channels is critical for hair-cell synapse function and can impede ribbon formation. We show that in mature zebrafish hair cells, evoked presynaptic-Ca^2+^ influx through Ca_V_1.3 channels initiates mitochondrial-Ca^2+^ (mito-Ca^2+^) uptake adjacent to ribbons. Block of mito-Ca^2+^ uptake in mature cells depresses presynaptic Ca^2+^ influx and impacts synapse integrity. In developing zebrafish hair cells, mito-Ca^2+^ uptake coincides with spontaneous rises in presynaptic Ca^2+^ influx. Spontaneous mito-Ca^2+^ loading lowers cellular NAD^+^/NADH redox and downregulates ribbon formation. Direct application of NAD^+^ or NADH increases or decreases ribbon formation respectively, possibly acting through the NAD(H)-binding domain on Ribeye. Our results present a mechanism where presynaptic- and mito-Ca^2+^ couple to confer proper presynaptic function and formation.

## Introduction

Neurotransmission is an energy demanding process that relies heavily on mitochondria. In neurons, mitochondria dysfunction has been implicated in synaptopathies that impact neurodevelopment, learning and memory, and can contribute to neurodegeneration (Flippo and Strack, 2017; Lepeta et al., 2016; Todorova and Blokland, 2017). In hair cells, sensory neurotransmission relies on specialized ribbon synapses to facilitate rapid and sustained vesicle release that is particularly energy demanding <u>(reviewed in: Johnson et al., 2019; Lagnado and Schmitz, 2015; Matthews and Fuchs, 2010; Safieddine et al., 2012)</u>. Although mitochondria dysfunction has been implicated in hearing loss (Böttger and Schacht, 2013; Fischel-Ghodsian et al., 2004; Kokotas et al., 2007), the precise role mitochondria play at hair-cell synapses remains unclear.

Ribbon synapses are characterized by a unique presynaptic structure called a “ribbon” that tethers and stabilizes synaptic vesicles at the active zone <u>(reviewed in: Matthews and</u> <u>Fuchs, 2010)</u>. In hair cells, neurotransmission at ribbon synapses requires the presynaptic-Ca^2+^ channel Ca_V_1.3 (Brandt et al., 2003; Kollmar et al., 1997; Sidi et al., 2004). Hair-cell depolarization opens Ca_V_1.3 channels, resulting in a spatially restricted increase of Ca^2+^ at presynaptic ribbons that triggers vesicle fusion. Tight spatial regulation of presynaptic Ca^2+^ is important for ribbon-synapse function and requires efficient Ca^2+^ clearance through a combination of Ca^2+^ pumps, Ca^2+^ buffers and intracellular Ca^2+^ stores (Carafoli, 2011; Mulkey and Malenka, 1992; Tucker and Fettiplace, 1995; Yamoah et al., 1998; Zenisek and Matthews, 2000). While ER Ca^2+^ stores have been implicated in hair-cell neurotransmission, whether mitochondrial-Ca^2+^ (mito-Ca^2+^) stores play a role in this process remains unclear (Castellano-Muñoz and Ricci, 2014; Kennedy, 2002; Lioudyno et al., 2004; Tucker and Fettiplace, 1995).

In addition to a role in hair-cell neurotransmission, presynaptic Ca^2+^ and Ca_V_1.3 channels also play an important role during inner-ear development. In mammals, prior to hearing onset, auditory hair cells fire spontaneous Ca^2+^ action potentials (Eckrich et al., 2018; Marcotti et al., 2003; Tritsch et al., 2007, 2010). In mammalian hair cells, these Ca^2+^ action potentials are Ca_V_1.3-dependent and are thought to be important for synapse and circuit formation. In support of this idea, *in vivo* work in zebrafish hair cells found that increasing or decreasing voltage-gated Ca^2+^ influx through Ca_V_1.3 channels during development led to the formation of smaller or larger ribbons respectively (Sheets et al., 2012). Furthermore, in mouse knockouts of Ca_V_1.3, auditory outer hair cells have reduced afferent innervation and synapse number (Ceriani et al., 2019). Mechanistically, how Ca_V_1.3-channel activity regulates ribbon size and innervation, and whether hair-cell Ca^2+^ stores play a role in this process is not known.

Cumulative work has shown that ribbon size varies between species and sensory epithelia <u>(reviewed in Moser et al., 2006</u>); these variations are thought to reflect important encoding requirements of a given sensory cell (Matthews and Fuchs, 2010). In auditory hair cells, excitotoxic noise damage can also alter ribbon size, and lead to hearing deficits (Jensen et al., 2015; Kujawa and Liberman, 2009; Liberman et al., 2015). Excitotoxic damage is thought to be initiated by mito-Ca^2+^ overload and subsequent ROS production (Böttger and Schacht, 2013; Wang et al., 2018). Mechanistically, precisely how ribbon size is established during development or altered under pathological conditions is not fully understood.

One known way to regulate ribbon formation is through its main structural component Ribeye (Schmitz et al., 2000a). Perhaps unsurprisingly, previous work has shown that overexpression or depletion of Ribeye in hair cells can increase or decrease ribbon size respectively (Becker et al., 2018; Jean et al., 2018; Sheets, 2017; Sheets et al., 2011a). Ribeye is a splice variant of the transcriptional co-repressor Carboxyl-terminal binding protein 2 (CtBP2) – a splice variant that is unique to vertebrate evolution (Schmitz et al., 2000a). Ribeye contains a unique A-domain, and a B-domain that is nearly identical to full-length CtBP2. The B-domain contains a nicotinamide adenine dinucleotide (NAD^+^, NADH or NAD(H)) binding site (Schmitz et al., 2000; Magupalli et al., 2008). NAD(H) redox is linked to mitochondrial metabolism (Srivastava, 2016). Because CtBP is able to bind and detect NAD^+^ and NADH levels, it is thought to function as a metabolic biosensor (Stankiewicz et al., 2014). For example, previous work has demonstrated that changes in NAD(H) redox can impact CtBP oligomerization and its transcriptional activity (Fjeld et al., 2003; Thio et al., 2004). Interestingly, *in vitro* work has shown that both NAD^+^ and NADH can also promote interactions between Ribeye domains (Magupalli et al., 2008). Whether NAD^+^ or NADH can impact Ribeye interactions and ribbon formation or stability has not been confirmed *in vivo*.

In neurons, it is well established that during presynaptic activity, mitochondria clear and store Ca^2+^ at the presynapse (Devine and Kittler, 2018). Additionally, presynaptic activity and mito-Ca^2+^ can couple together to influence cellular bioenergetics, including NAD(H) redox homeostasis (reviewed in: <u>Kann and Kovács, 2007; Llorente-Folch et al., 2015)</u>. Based on these studies, we hypothesized that Ca^2+^ influx through Ca_V_1.3 channels may regulate mito-Ca^2+^, which in turn could regulate NAD(H) redox. Changes to cellular bioenergetics and NAD(H) redox could function to control Ribeye interactions and ribbon formation or impact ribbon-synapse function and stability.

To study the impact of mito-Ca^2+^ and NAD(H) redox on ribbon synapses, we examined hair cells in the lateral-line system of larval zebrafish. This system is advantageous for our studies because it contains hair cells with easy access for *in vivo* pharmacology, mechanical stimulation and imaging cellular morphology and function. Within the lateral-line, hair cells are arranged in clusters called neuromasts. The hair cells and ribbon synapses in each cluster form rapidly between 2 to 3 days post-fertilization (dpf) but by 5-6 dpf, the majority of hair cells are mature, and the system is functional (Kindt et al., 2012; McHenry et al., 2009; Metcalfe, 1985; Murakami et al., 2003; Santos et al., 2006). Thus, these two ages (2-3 dpf and 5-6 dpf) can be used to study mito-Ca^2+^ and NAD(H) redox in developing and mature hair cells respectively.

Using this sensory system, we find that presynaptic Ca^2+^ influx drives mito-Ca^2+^ uptake. In mature hair cells, mito-Ca^2+^ uptake occurs during evoked stimulation and is required to sustain presynaptic function and ultimately synapse integrity. In developing hair cells, mito-Ca^2+^ uptake coincides with spontaneous rises in presynaptic Ca^2+^. Blocking these spontaneous changes in Ca^2+^ leads to the formation of larger ribbons. Using a redox biosensor, we demonstrate that specifically in developing hair cells, decreasing mito-Ca^2+^ levels increases the NAD^+^/NADH redox ratio. Furthermore, we show that application of NAD^+^ or NADH can increase or decrease ribbon formation respectively. Overall our results suggest that in hair cells presynaptic Ca^2+^ influx and mito-Ca^2+^ uptake couple to impact ribbon formation and function.

## Results

### Mitochondria are located near presynaptic ribbons

In neurons, synaptic mitochondria have been shown to influence synapse formation, plasticity and function (Flippo and Strack, 2017; Todorova and Blokland, 2017). Based on this work, we hypothesized that mitochondria may impact synapses in hair cells. Therefore, we examined the proximity of mitochondria relative to presynaptic ribbons in zebrafish lateral-line hair cells. We visualized mitochondria and ribbons using transmission electron microscopy (TEM) and in live hair cells using Airyscan confocal microscopy.

Using TEM, we examined sections that clearly captured ribbons (Example, Figure 1C). We were able to observe a mitochondrion in close proximity (< 1 µm) to ribbons in 74 % of the sections (Figure 1D, median ribbon-to-mitochondria distance = 174 nm, n = 17 out of 21 sections). To obtain a more comprehensive understanding of the 3D morphology and location of mitochondria relative to ribbons in live cells, we used Airyscan confocal microscopy. To visualize these structures in living cells, we used transgenic zebrafish expressing MitoGCaMP3 (Esterberg et al., 2014) and Ribeye a-tagRFP (Sheets et al., 2017) in hair cells to visualize mitochondria and ribbons respectively. Using this approach, we observed tubular networks of mitochondria extending from apex to base (Figure 1A-B, E-E’, Figure S1A, Movie S1). At the base of the hair cell, we observed ribbons nestled between branches of mitochondria. Overall our TEM and Airyscan imaging suggests that in lateral-line hair cells, mitochondria are present near ribbons and are poised to impact ribbon synapses.

**Figure 1.**
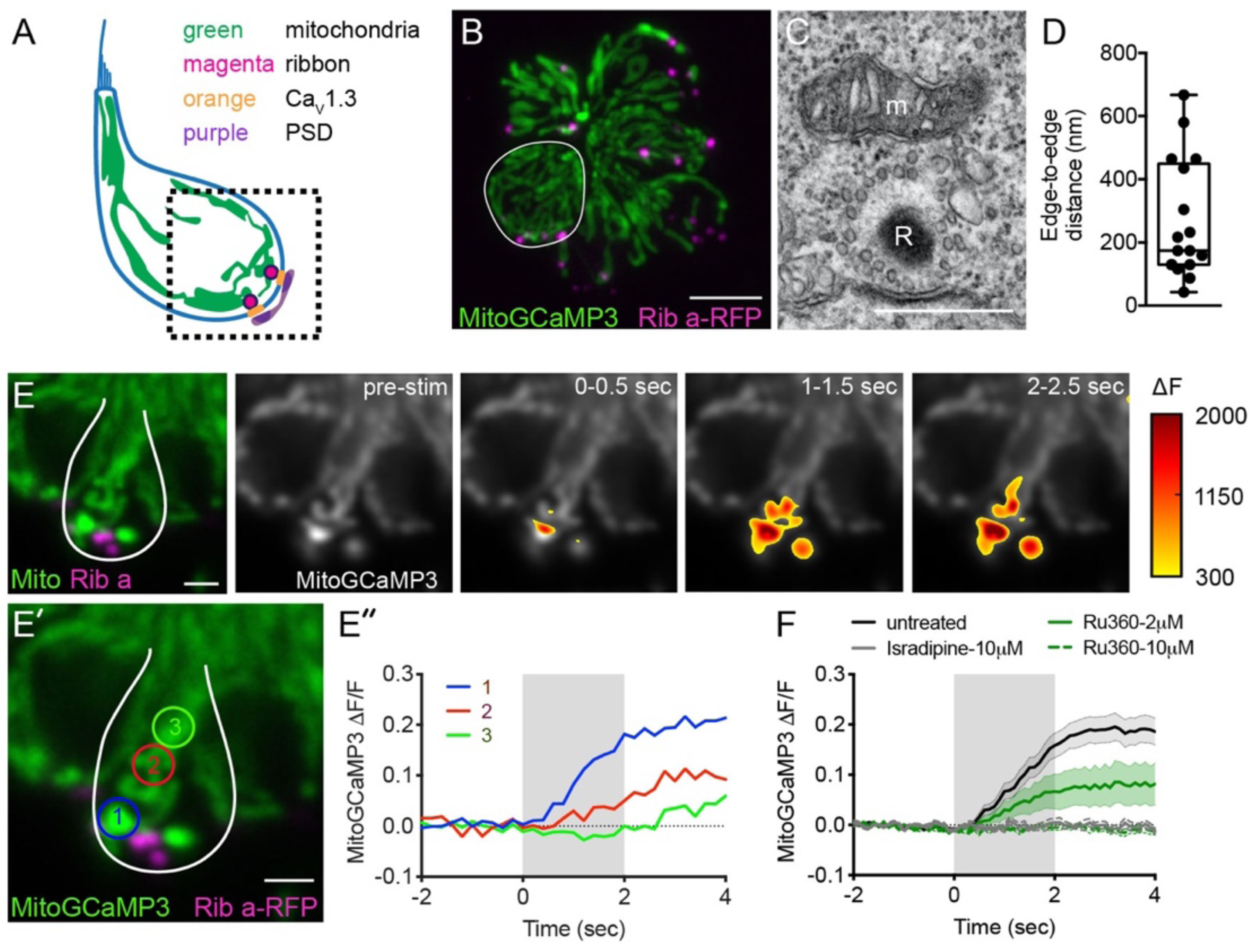
Mito-Ca^2+^ uptake initiates adjacent to ribbons. A, cartoon illustration of a lateral-line hair cell containing: an apical mechanosensory bundle (blue), mitochondria (green), presynaptic ribbons (magenta), Ca_V_1.3 channels (orange) and postsynaptic densities (purple). B, Airyscan confocal image of 6 live hair cells (1 cell outlined in white) expressing MitoGCaMP3 (mitochondria) and Ribeye a-tagRFP (ribbons) in a developing neuromast at 2 dpf. Also see Figure S1. C, A representative TEM showing a mitochondrion (m) in close proximity to a ribbon (R) at 4 dpf. D, Quantification of mitochondrion to ribbon distance in TEM sections (n = 17 sections). E, Side-view of a hair cell (outlined in white) shows the spatio-temporal dynamics of evoked mito-Ca^2+^ signals during a 2-s stimulation at 6 dpf. The MitoGCaMP3 signals are indicated by the heatmap and are overlaid onto the pre-stimulus grayscale image. E’-E’’, Circles 1-3 (1.3 μm diameter) denote regions used to generate the temporal traces of mito-Ca^2+^ signals in E’’: adjacent to the presynapse (“1”), and midbody (“2” and “3”) in the same cell as E. F, Average evoked mito-Ca^2+^ response before (solid black) and after 30 min incubation with 10 μM Ru360 (dashed green), 2 μM Ru360 (green), or 10 μM isradipine (gray) (3-5 dpf, n ≥ 9 cells per treatment). Error bars in D are min and max; in F the shaded area denotes SEM. Scale bar = 500 nm in C, 5 µm in B and 2 µm in E and E’.

### Mito-Ca^2+^ uptake at ribbons is MCU and Ca_V_1.3 dependent

In zebrafish hair cells, robust rises in mito-Ca^2+^ have be reported during mechanical stimulation (Pickett et al., 2018). Due to the proximity of the mitochondria to the ribbon, we predicted that rises in mito-Ca^2+^ levels during mechanical stimulation are related to presynapse-associated rises in Ca^2+^.

To test this prediction, we used a fluid-jet to mechanically stimulate hair cells and evoke presynaptic activity. During stimulation, we used MitoGCaMP3 to monitor mito-Ca^2+^ in hair cells. As previously reported, we observed robust mito-Ca^2+^ uptake during stimulation (Figure 1E-F). We examined the subcellular distribution of MitoGCaMP3 signals over time and found that the signals initiated near ribbons (Figure 1E). During the latter part of the stimulus, and even after the stimulus terminated, the MitoGCaMP3 signals propagated apically within the mitochondria, away from the ribbons (Example, Figure 1E-E’’, regions 1-3). We characterized the time course of MitoGCaMP3 signals with regards to onset kinetics and return to baseline. During a 2-s stimulus, we detected a significant rise in MitoGCaMP3 signals 0.6 s after stimulus onset (Figure S1B). Interestingly, after the stimulus terminated, MitoGCaMP3 levels took approximately 5 min to return to baseline (Figure S1C-C’). As previously reported, the kinetics of MitoGCaMP3 signals in hair cells mitochondria were quite different from signals observed using cytosolic GCaMP3 (CytoGCaMP3) in hair cells (Pickett et al., 2018). Compared to MitoGCaMP3 signals, CytoGCaMP3 signals had faster onset kinetics, and a faster return to baseline (Figure S1B-C, time to rise: 0.06 s, post-stimulus return to baseline: 12 s). These differences in kinetics indicate that mito-Ca^2+^ loading operates over slower timescales compared to the cytosolic compartment. It also confirms that hair-cell stimulation can initiate long lasting increases in mito-Ca^2+^.

To verify that MitoGCaMP3 signals reflect Ca^2+^ entry into mitochondria, we applied Ru360, an antagonist of the mito-Ca^2+^ uniporter (MCU). The MCU is the main pathway for rapid Ca^2+^ entry into the mitochondria (Matlib et al., 1998). We found that stimulus-evoked MitoGCaMP3 signals were blocked in a dose-dependent manner after treatment with Ru360 (Figure 1F). Due to the initiation of mito-Ca^2+^ near ribbons, we examined whether presynaptic Ca^2+^ influx through Ca_V_1.3 channels was the main source of Ca^2+^ entering the mitochondria. To examine Ca_V_1.3 channel contribution to mito-Ca^2+^ uptake, we applied isradipine, a Ca_V_1.3 channel antagonist. Similar to blocking the MCU, blocking Ca_V_1.3 channels eliminated all stimulus-evoked MitoGCaMP3 signals (Figure 1F). Overall our MitoGCaMP3 functional imaging indicates that in hair cells, evoked mito-Ca^2+^ uptake initiates near ribbons and is dependent on MCU and Ca_V_1.3 channel function.

### Mito-Ca^2+^ uptake occurs in cells with presynaptic Ca^2+^ influx

Interestingly, we observed that mito-Ca^2+^ uptake was only present in ∼40 % of cells (Example, Figure 2A’; n = 10 neuromasts, 146 cells). This observation is consistent with previous work demonstrating that only ∼30 % of hair cells within each neuromast cluster have presynaptic Ca^2+^ signals and are synaptically active (Zhang et al., 2018b). Because presynaptic Ca^2+^ signals initiate near mitochondria, it is probable that mito-Ca^2+^ uptake may occur specifically in hair cells with synaptic activity.

**Figure 2.**
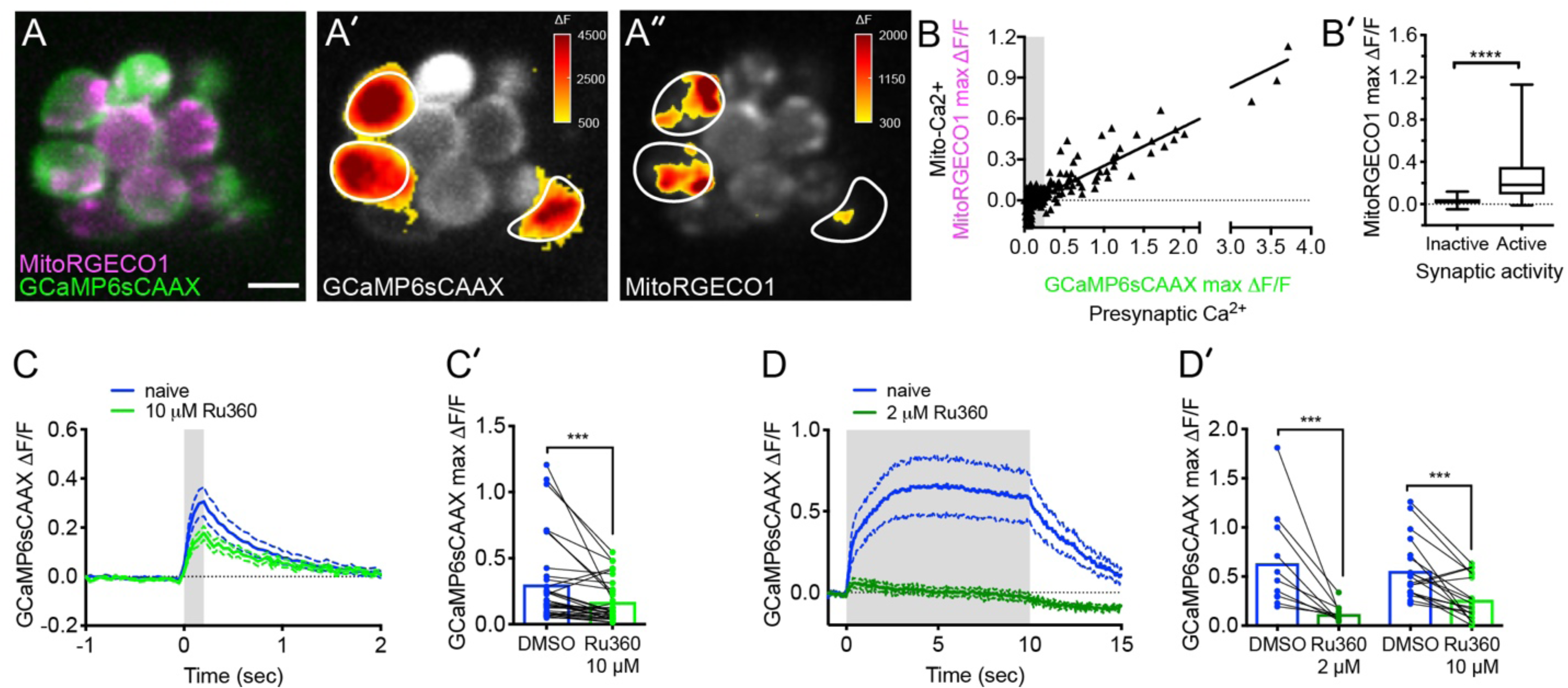
Mito-Ca^2+^ uptake can impact presynaptic Ca^2+^ signals. A, A live Image of a neuromast viewed top-down, expressing the presynaptic-Ca^2+^ sensor GCaMP6sCAAX (green) and mito-Ca^2+^ sensor MitoRGECO1 (magenta) at 3 dpf. The GCaMP6sCAAX (A’) and MitoRGECO1 (A’’) signals during a 2-s stimulation are indicated by the heatmaps and occur in the same cells (white outline). B, Scatterplot with linear regression of peak presynaptic- and mito-Ca^2+^ response for individual cells at 3-5 dpf, n = 209 cells. Gray background in graph denotes presynaptic-Ca^2+^ signals below 0.25, a threshold used as a cutoff for presynaptic activity (below inactive, above active). B’, Plot of mito-Ca^2+^responses segregated based on the activity threshold in B. C-D’, Presynaptic-Ca^2+^ response (example in Figure S2) averaged per cell before (blue) and after 30 min of 10 μM Ru360 (light green) or 2 μM Ru360 (dark green), n ≥ 10 cells per treatment. C and D show averaged traces while C’ and D’ show before-and-after dot plots of the peak response per cell. Whiskers on plots in B’ represent min and max; error (dashed lines) in plots C and D represent SEM. Mann-Whitney U test was used in B’; Wilcoxon matched-pairs signed-rank test was used in C’ and D’. \*\*\**p* < 0.001, \*\*\*\**p* < 0.0001. Scale bar = 5 µm in A.

To test whether evoked mito-Ca^2+^ uptake occurred exclusively in cells with presynaptic Ca^2+^ influx, we performed two-color functional imaging. We used a double transgenic approach that utilized a membrane-localized GCaMP6s (GCaMP6sCAAX; green) to measure presynaptic Ca^2+^ signals at the base of hair cells (Jiang et al., 2017a; Sheets et al., 2017), and concurrently used MitoRGECO1 (red) to examine mito-Ca^2+^ signals (Figure 2A-B’). Our two-color imaging approach revealed a strong correlation between the magnitude of the GCaMP6sCAAX and MitoRGECO1 signals (Figure 2B, R^2^ = 0.8, p < 0.0001; n = 209 cells). We found that the median MitoRGECO1 signals were 400 % larger in presynaptically active hair cells compared to presynaptically silent hair cells (Figure 2B’). Together these results suggest that mito-Ca^2+^ uptake occurs specifically in hair cells with evoked presynaptic-Ca^2+^ influx.

### Blocking Mito-Ca^2+^ entry depresses presynaptic Ca^2+^ signals in mature hair cells

Although we observed mito-Ca^2+^ uptake specifically in hair cells with active Ca^2+^ channels, the impact of mito-Ca^2+^ uptake on the function of hair-cell synapses was unclear. Based on previous studies in neurons (Billups and Forsythe, 2002; Levy et al., 2003; Chouhan et al., 2010; Kwon et al., 2016), we reasoned that mitochondria may also be important to remove excess Ca^2+^ from the hair-cell presynapse to regulate neurotransmission.

To determine if mito-Ca^2+^ uptake impacted presynaptic function, we assayed evoked presynaptic-Ca^2+^ signals by monitoring GCaMP6sCAAX signals adjacent to ribbons as described previously (Example, Figure S2, Sheets et al., 2017; Zhang et al., 2018b). We examined GCaMP6sCAAX signals in mature-hair cells at 5-6 dpf when neuromast organs are largely mature (Kindt et al., 2012; McHenry et al., 2009; Metcalfe, 1985; Murakami et al., 2003; Santos et al., 2006). Using this approach, we assayed presynaptic GCaMP6sCAAX signals before and after a 20-min application of the MCU antagonist Ru360 (Figure 2C-D’). We found that during short, 200-ms stimuli, GCaMP6sCAAX signals at ribbons were reduced after complete MCU block (10 µM Ru360, Figure 2C-C’). Reduction of GCaMP6sCAAX signals were further exacerbated during sustained 10-s stimuli, even when the MCU was only partially blocked (2 µM Ru360, Figure 2D-D’). These results suggest that in mature hair cells, evoked mito-Ca^2+^ uptake is critical for presynaptic Ca^2+^ influx, especially during sustained stimulation.

### Evoked mito-Ca^2+^ uptake is important for mature synapse integrity and cell health

MCU block could impair presynaptic Ca^2+^ influx through several mechanisms. It could impair the biophysical properties of Ca_V_1.3 channels, for example, through Ca^2+^-dependent inactivation (Platzer et al., 2000; Schnee and Ricci, 2003). In addition, mito-Ca^2+^ has been implicated in synapse dysfunction and cell death (Esterberg et al., 2014; Vos et al., 2010; Wang et al., 2018), and MCU block could be pathological. To distinguish between these possibilities, we assessed whether synapse or hair-cell number were altered after MCU block with Ru360.

To quantify ribbon-synapse morphology after MCU block, we immunostained mature-hair cells (5 dpf) with Ribeye b and MAGUK antibodies to label presynaptic ribbons and postsynaptic densities (MAGUK) respectively. We first applied 2 μM Ru360 for 1 hr, a concentration that partially reduces evoked mito-Ca^2+^ uptake (See Figure 1F’) yet is effective at reducing sustained presynaptic Ca^2+^ influx (See Figure 2D-D’). At this dose, Ru360 had no impact on hair cell or synapse number (Figure 3E). In addition, we observed no morphological change in ribbon or postsynapse size (Figure 3F, Figure S3A). These findings indicate that partial MCU block can impair presynaptic function without any observable pathology.

**Figure 3.**
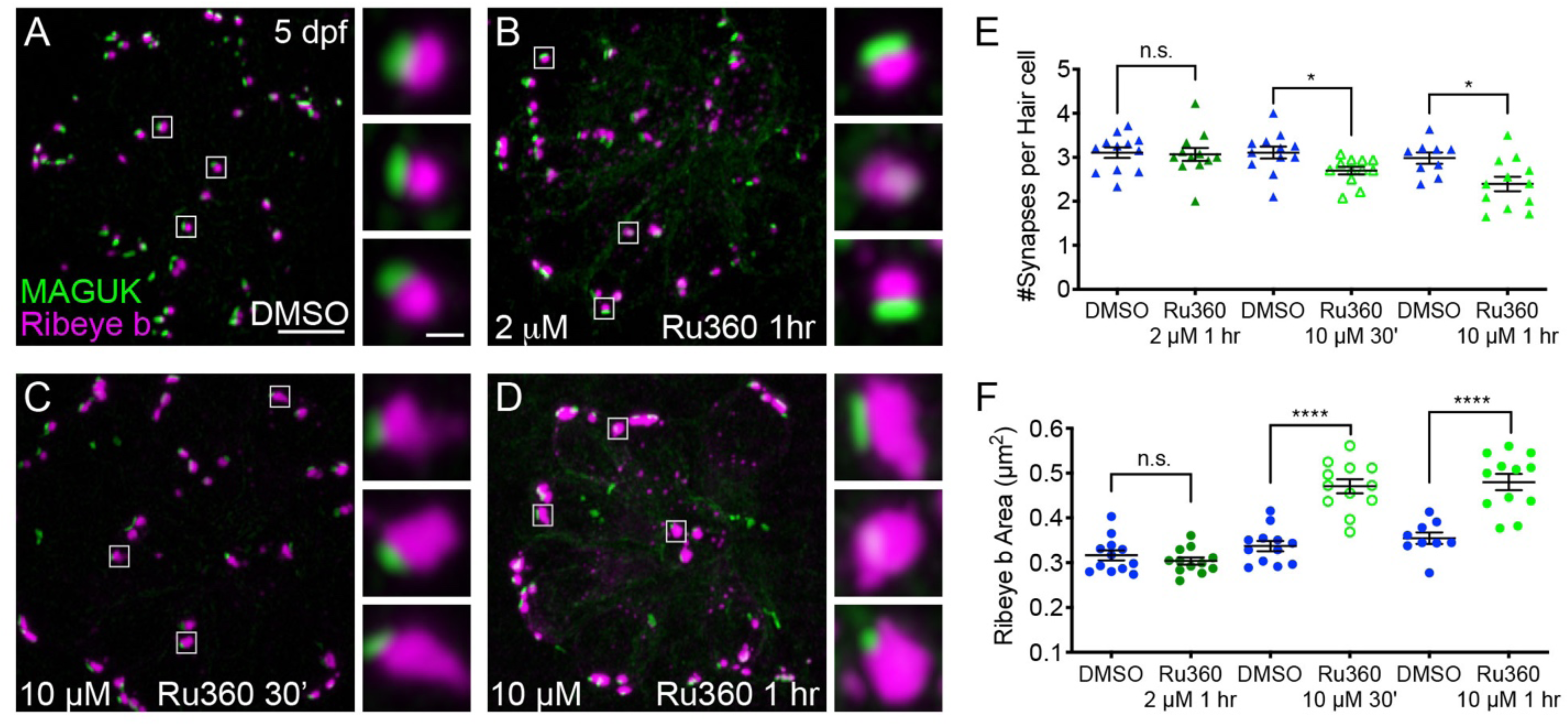
Mito-Ca^2+^ is important for ribbon size and synapse integrity in mature hair cells. A-D, Representative images of mature neuromasts (5 dpf) immunostained with Ribeye b (magenta, ribbons) and MAGUK (green, postsynapses) after a 1 hr 0.1% DMSO (A), a 1 hr 2 μM Ru360 (B), a 30 min 10 μM Ru360 (C), or a 1 hr 10 μM Ru360 (D) treatment. Insets show 3 example synapses (white squares). E-F, Scatter plots show synapse counts (E), and ribbon area (F) in controls and in treatment groups. N ≥ 9 neuromasts per treatment. Error bars in E-F represent SEM. A Welch’s unequal variance *t*-test was used in E-F. \**p* < 0.05, \*\*\*\**p* < 0.0001. Scale bar = 5 µm in A, and 2 µm in inset.

We also tested a higher dose of Ru360 (10 µM) that completely blocks evoked mito-Ca^2+^ uptake (See Figure 1F). Interestingly, a 30-min or 1-hr 10 µM Ru360 treatment had a progressive impact on synapse and cellular integrity. After a 30-min treatment with 10 µM Ru360 we observed significantly fewer complete synapses per hair cell, but not fewer hair cells compared to controls (Figure 3E; Hair cells per neuromast, control: 16.3, 30-min 10 µM Ru360: 15.5; p = 0.5). In addition, after the 30-min treatment, ribbons were significantly larger (Figure 3F). The pathological effects of MCU block were more pronounced after a 1-hr, 10 µM Ru360 treatment. After 1-hr, there was both fewer hair cells per neuromast (Hair cells per neuromast, control: 18.1, 1-hr 10 µM Ru360: 12.0; p > 0.0001) and fewer synapses per hair cell (Figure 3E). Similar to 30-min treatments with Ru360, after 1 hr, ribbons were also significantly larger (Figure 3F). Neither 30-min nor 1-hr 10 µM Ru360 treatment altered postsynapse size (Figure S3A). Overall, our results indicate that in mature hair cells, partial block of mito-Ca^2+^ uptake can impair presynaptic function without altering presynaptic morphology or synapse integrity. Complete block of mito-Ca^2+^ uptake is pathological; it impairs presynaptic function, alters presynaptic morphology, and results in a loss of synapses and hair-cells.

### Spontaneous presynaptic and mito-Ca^2+^ influx pair in developing hair cells

In addition to evoked presynaptic- and mito-Ca^2+^ signals in hair cells, we also observed instances of spontaneous presynaptic- and mito-Ca^2+^ signals (Example, Figure 4A-A’’’, Movie S2). Numerous studies have demonstrated that mammalian hair cells have spontaneous presynaptic-Ca^2+^ influx during development (Eckrich et al., 2018; Marcotti et al., 2003; Tritsch et al., 2007, 2010). Therefore, we predicted that similar to mammals, spontaneous presynaptic- Ca^2+^ uptake may be a feature of development. Furthermore, we predicted that spontaneous mito-Ca^2+^ uptake may correlate with instances of spontaneous presynaptic-Ca^2+^ influx.

**Figure 4.**
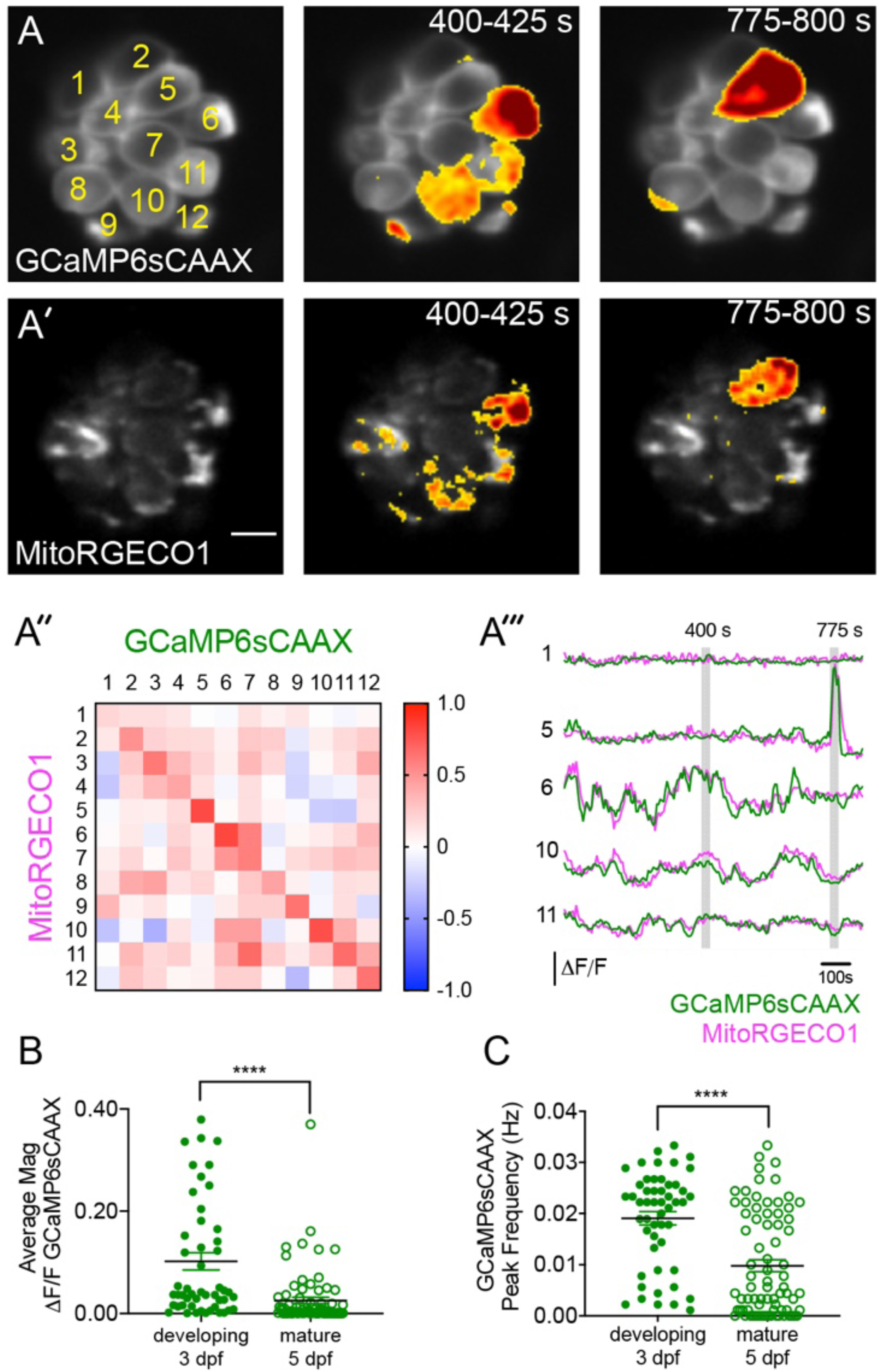
Spontaneous presynaptic- Ca^2+^ influx and Mito-Ca^2+^ uptake are linked. A-A’, A live Image of a neuromast viewed top-down, expressing the presynaptic-Ca^2+^ sensor GCaMP6sCAAX (A) and mito-Ca^2+^ sensor MitoRGECO1 (A’) at 3 dpf. Example GCaMP6sCAAX (A’) and MitoRGECO1 (A’) signals during two 25-s windows within a 900-s acquisition are indicated by the ΔF heatmaps and occur in the same cells. A’’, A heatmap of Pearson correlation coefficients comparing GCaMP6sCAAX and MitoRGECO1 signals from the cells in A-A’. A’’’, Example GCaMP6sCAAX (green) MitoRGECO1 (magenta) traces during the 900-s acquisition from the 5 cells numbered in A, also see Movie S2. B, Scatterplot showing the average magnitude of GCaMP6sCAAX signals in developing and mature hair cells, n = 6 neuromasts per age. C, Scatterplot showing frequency of GCaMP6sCAAX events in developing and mature hair cells, n = 6 neuromasts. Error bars in B-C represent SEM. A Mann-Whitney U test was used in B and C. \*\*\*\**p* < 0.0001. Scale bar = 5 µm in A.

First we tested whether spontaneous presynaptic-Ca^2+^ signals were a feature of development. In zebrafish neuromasts, hair cells are rapidly added between 2-3 dpf, but by 5-6 dpf relatively fewer cells are added and the hair cells and the organs are largely mature (Kindt et al., 2012; McHenry et al., 2009; Metcalfe, 1985; Murakami et al., 2003; Santos et al., 2006). Therefore, we examined the magnitude and frequency of spontaneous, presynaptic GCaMP6sCAAX signals in developing (3 dpf) and mature hair cells (5 dpf). We found that in developing hair cells, spontaneous GCaMP6sCAAX signals occurred with larger magnitudes and more frequency compared to those in mature hair cells (Figure 4B-C). Our spontaneous GCaMP6sCAAX imaging demonstrates that similar to mammals, spontaneous presynaptic Ca^2+^ activity is a feature of developing zebrafish hair cells.

Next, we tested whether spontaneous mito-Ca^2+^ uptake and presynaptic-Ca^2+^ influx were correlated. For this analysis we concurrently imaged GCaMP6sCAAX and MitoRGECO1 signals in the same cells for 15 mins to measure presynaptic and mito-Ca^2+^ responses respectively. We found that spontaneous presynaptic-Ca^2+^ influx was often associated with spontaneous mito-Ca^2+^ uptake (Example, Figure 4A-A’’’). Overall, we observed a high correlation between the rise and fall of these two signals within individual cells (Figure A’’-A’’’). Both of these signals and their correlation are abolished by application of the Ca_V_1.3-channel antagonist isradipine (Figure S4). Together these experiments indicate that, similar to our evoked experiments, spontaneous presynaptic- and mito- Ca^2+^ signals are correlated.

### Spontaneous mito-Ca^2+^ uptake regulates ribbon formation

Previous work in zebrafish demonstrated that Ca_V_1.3 channel activity plays a role in ribbon formation specifically during development (Sheets et al., 2012). This work found that a transient, 1-hr pharmacological block of Ca_V_1.3 channels increased ribbon size, while Ca_V_1.3 channel agonists decreased ribbon size (Figure 5E; Sheets et al., 2012). Therefore, spontaneous Ca_V_1.3 and MCU Ca^2+^ activities could function together to control ribbon size in developing hair cells.

**Figure 5.**
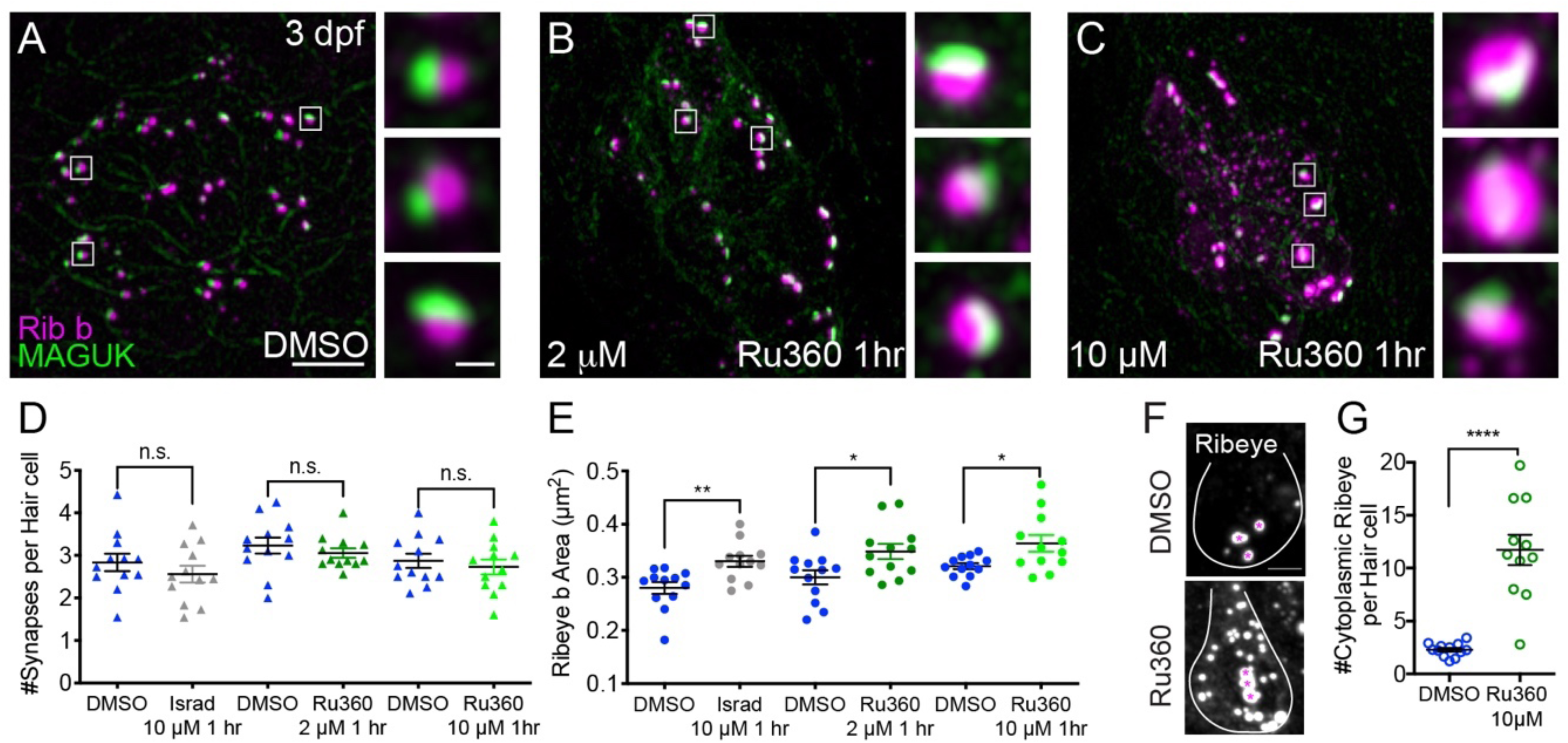
Mito-Ca^2+^ regulates ribbon formation. A-C, Representative images of immature neuromasts (3 dpf) immunostained with Ribeye b (magenta, ribbons) and MAGUK (green, postsynapses) after a 1 hr 0.1% DMSO (A), 2 μM Ru360 (B) or 10 µM Ru360 (C) treatment. Insets show 3 representative synapses (white squares) for each treatment. (D-E) Scatterplot show quantification of synapse number (D), and ribbon area (E) in controls and in treatment groups. F, Side-view of hair cell (white outline) shows synaptic ribbon (magenta asterisks) and extrasynaptic Ribeye b aggregates after a 1 hr 0.1% DMSO or 10 μM Ru360 treatment. Quantification of extrasynaptic Ribeye puncta (G). N ≥ 12 neuromasts per treatment. Error bars in B-C represent SEM. Welch’s unequal variance *t*-test was used in D-E and G, \**p* < 0.05, \*\**p* < 0.01, \*\*\*\**p*<0.0001. Scale bar = 5 µm in A, 2 µm in insets and F.

To characterize the role of MCU function and spontaneous mito-Ca^2+^ uptake on ribbon formation, we applied the MCU antagonist Ru360 to developing hair cells (3 dpf). After this treatment, we quantified ribbon synapse morphology by immunostaining hair cells to label presynaptic ribbons and postsynaptic densities. After a 1-hr application of 2 μM Ru360 to block the MCU, we observed a significant increase in ribbon size in developing hair cells (Figure 5A-B, E). In contrast, this same treatment did not impact ribbon size in mature hair cells (Figure 3F). We also applied a higher concentration of Ru360 (10 µM) to developing hair cells for 1 hr. In developing hair cells, after a 1-hr 10 µM Ru360 treatment, we also observed a significant increase in ribbon size (Figure 5A, C, E). Unlike in mature hair cells (Figure 3), in developing hair cells, these concentrations of the MCU antagonist did not alter the number of hair cells, nor the number of synapses per hair cell (Figure 5D; Hair cells per neuromast, control: 9.0, 1-hr 10 µM Ru360: 8.8, p = 0.3). All morphological changes were restricted to the ribbons, as MCU block did not alter the size of the postsynapse (Figure S3C).

In addition to larger ribbons, at higher concentrations of Ru360 (10 µM) we also observed an increase in cytoplasmic, non-synaptic Ribeye aggregates (Figure 5F, G). Previous work in zebrafish reported both larger ribbons and cytoplasmic aggregates of Ribeye in Ca_V_1.3a-deficient hair cells (Sheets et al., 2011a). These parallel phenotypes indicate that spontaneous presynaptic Ca^2+^ influx and mito-Ca^2+^ uptake may couple to shape ribbon formation. Our results suggest that during development, spontaneous Ca^2+^ entry through both Ca_V_1.3 and MCU channels continuously regulate ribbon formation; blocking either channel increases Ribeye aggregation and ribbon size.

### MCU and Ca_V_1.3 channel activities regulate subcellular Ca^2+^ homeostasis

Our results indicate that spontaneous Ca^2+^ influx through Ca_V_1.3 channels and subsequent loading of Ca^2+^ into mitochondria regulates ribbon formation in developing hair cells. But how do these two Ca^2+^ signals converge to regulate ribbon formation? It is possible that mitochondria could buffer Ca^2+^ during spontaneous presynaptic activity and function to decrease resting levels of cytosolic Ca^2+^ (cyto-Ca^2+^); cyto-Ca^2+^ levels could be a signal that regulates ribbon formation. To examine resting cyto-Ca^2+^ levels in hair cells, we examined the fluorescence signal change of the cytosolic Ca^2+^ indicator RGECO1 (CytoRGECO1) before and after a 30-min pharmacological manipulation of Ca_V_1.3 or MCU channels (Figure 6A-C).

**Figure 6.**
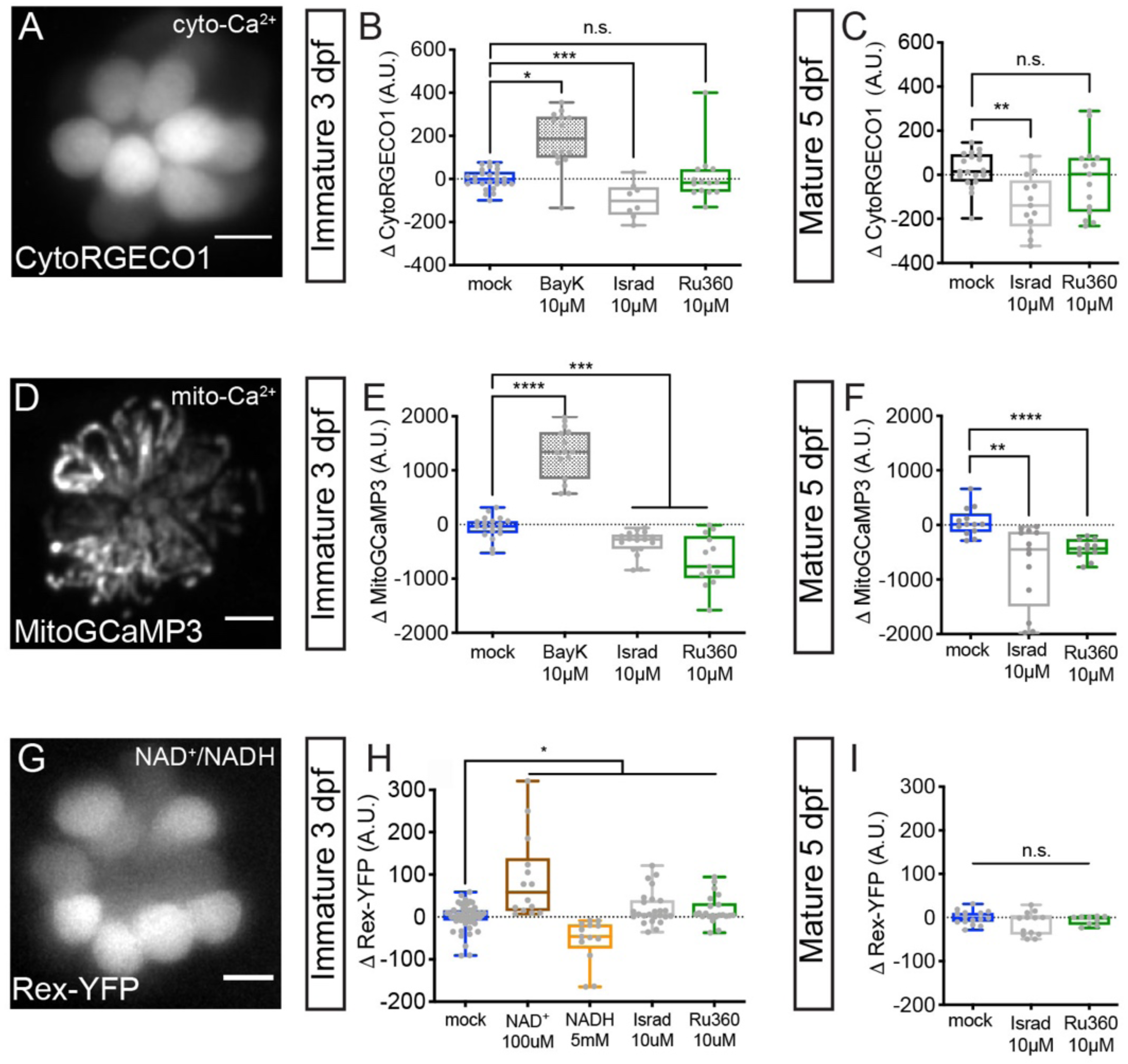
Cyto-Ca^2+^, mito-Ca^2+^ and NAD^+^/NADH redox baseline measurements. Live hair cells expressing RGECO1 (A), MitoGCaMP3 (D), or Rex-YFP (G) show resting cyto-Ca^2+^, mito-Ca^2+^ or NAD^+^/NADH levels respectively. B-C, RGECO1 baseline measurements before and after a 30 min mock treatment (0.1% DMSO), or after a 30 min 10 μM Bay K8644 (BayK), 10 μM isradipine, or 10 μM Ru360 treatment. E-F, MitoGCaMP3 baseline measurements before and after a 30 min mock treatment (0.1% DMSO), or after a 10 μM BayK, 10 μM isradipine, or 10 μM Ru360 treatment. H-I, Rex-YFP baseline measurements before and after 30 min mock treatment (0.1% DMSO), or after a 30 min 100 μM NAD^+^, 5 mM NADH, 10 μM isradipine, or 10 μM Ru360 treatment. All plots are box-and-whiskers plot that show median, min and max. N ≥ 9 neuromasts per treatment. One-way Brown-Forsythe and Welch ANOVA with Dunnett’s T3 post hoc was used to calculate the difference in B-C, E-F, and H-I, \**p* < 0.05, \*\**p* < 0.01, \*\*\**p* < 0.001, \*\*\*\**p* < 0.0001. Scale bar = 5 μm in A, D and G.

We observed that treatment with the Ca_V_1.3 channel antagonist isradipine and agonist Bay K8644 decreased and increased resting CytoRGECO1 fluorescence respectively (Figure 6B). However, treatment with MCU blocker Ru360 did not significantly shift resting CytoRGECO1 fluorescence levels (Figure 6B). Similar results with Ru360 were observed in developing and mature hair cells (Figure 6B-C). These data suggest that, unlike Ca_V_1.3 channel function, MCU function and associated mito-Ca^2+^ uptake does not play a critical role in buffering steady state cyto-Ca^2+^ levels.

Alternatively, it is possible that rather than impacting cyto-Ca^2+^ levels, both Ca_V_1.3 and MCU activity are required to load and maintain Ca^2+^ levels within the mitochondria. In this scenario, mito-Ca^2+^ levels could be a signal that regulates ribbon formation. To test this possibility, we used MitoGCaMP3 to examine resting mito-Ca^2+^ levels before and after modulating Ca_V_1.3 or MCU channel function (Figure 6D-F). We observed that blocking Ca_V_1.3 channels with isradipine or the MCU with Ru360 decreased resting MitoGCaMP3 fluorescence (Figure 6E-F). Conversely, Ca_V_1.3 channel agonist Bay K8644 increased resting MitoGCaMP3 fluorescence (Figure 6E). These results were consistent in developing and mature hair cells (Figure 6E-F). Our resting MitoGCaMP3 measurements indicate that the effects of Ca_V_1.3 channel and MCU activity converge in to regulate mito-Ca^2+^ levels. When either of these channels are blocked, the resting levels of mito-Ca^2+^ are decreased. Therefore, if presynaptic Ca^2+^ influx and mito-Ca^2+^ regulate ribbon formation through a similar mechanism, they may act through mito-rather than cyto-Ca^2+^ homeostasis.

### Mito-Ca^2+^ levels regulate NAD(H) redox in developing hair cells

If mito-Ca^2+^ levels signal to regulate ribbon formation, how is this signal transmitted from the mitochondria to the ribbon? An ideal candidate is via NAD(H) homeostasis. Ribeye protein, the main component of ribbons contains a putative NAD(H) binding site. Because mitochondria regulate NAD(H) redox homeostasis (Jensen-Smith et al., 2012) we reasoned that there may be a relationship between mito-Ca^2+^ levels, NAD(H) redox and ribbon formation.

To examine NAD(H) redox, we created a stable transgenic line expressing Rex-YFP, a fluorescent NAD^+^/NADH ratio biosensor in hair cells (Figure 6G). We verified the function of the Rex-YFP biosensor in our *in vivo* system by exogenously applying NAD^+^ or NADH for 30 min. We found that incubations with 100 µM NAD^+^ increased while 5 mM NADH decreased Rex-YFP fluorescence; these intensity changes are consistent with an increase and decrease in the NAD^+^/NADH ratio respectively (Figure 6H). Next, we examined if Ca_V_1.3 and MCU channel activities impact the NAD^+^/NADH ratio. We found that 30-min treatments with either Ca_V_1.3 or MCU channel antagonist increased the NAD^+^/NADH ratio (increased Rex-YFP fluorescence) in developing hair cells (Figure 6H). Interestingly, similar 30-min treatments did not alter Rex-YFP fluorescence in mature hair cells (Figure 6I). Together, our baseline MitoGCaMP3 and Rex-YFP measurements indicate that during development, Ca_V_1.3 and MCU channel activities normally function to increase mito-Ca^2+^ and decrease the NAD^+^/NADH ratio. Overall, this work provides strong evidence that links NAD(H) redox and mito-Ca^2+^ with ribbon formation.

### NAD^+^ and NADH directly influence ribbon formation

Our Rex-YFP measurements suggest that spontaneous Ca_V_1.3 and MCU Ca^2+^ activities normally function to decrease the NAD^+^/NADH ratio; furthermore, this activity may function to restrict ribbon formation. Conversely, blocking these activities increases the NAD^+^/NADH ratio and may increase ribbon formation. If the NAD^+^/NADH ratio is an intermediate step between Ca_V_1.3 and MCU channel activities and ribbon formation, we predicted that more NAD^+^ or NADH would increase or decrease ribbon formation respectively. To test this prediction, we treated developing hair cells with exogenous NAD^+^ or NADH.

After a 1-hr treatment with 100 µM NAD^+^, we found that the ribbons in developing hair cells were significantly larger compared to controls (Figure 7A-B, E). In contrast, after a 1-hr treatment with 5 mM NADH, ribbons were significantly smaller compared to controls (Figure 7A, C, E). Neither exogenous NAD^+^ nor NADH were able to alter ribbon size in mature hair cells (Figure 7F-H, J). These concentrations of NAD^+^ and NADH altered neither the number of synapses per hair cell nor postsynapse size in developing or mature hair cells (Figure 3D, I, Figure S3B, D). These results suggest that in developing hair cells, NAD^+^ promotes while NADH inhibits Ribeye-Ribeye interactions or Ribeye localization to the ribbon. Overall these results support the idea that during development, the levels of NAD^+^ and NADH can directly regulate ribbon formation *in vivo*.

**Figure 7.**
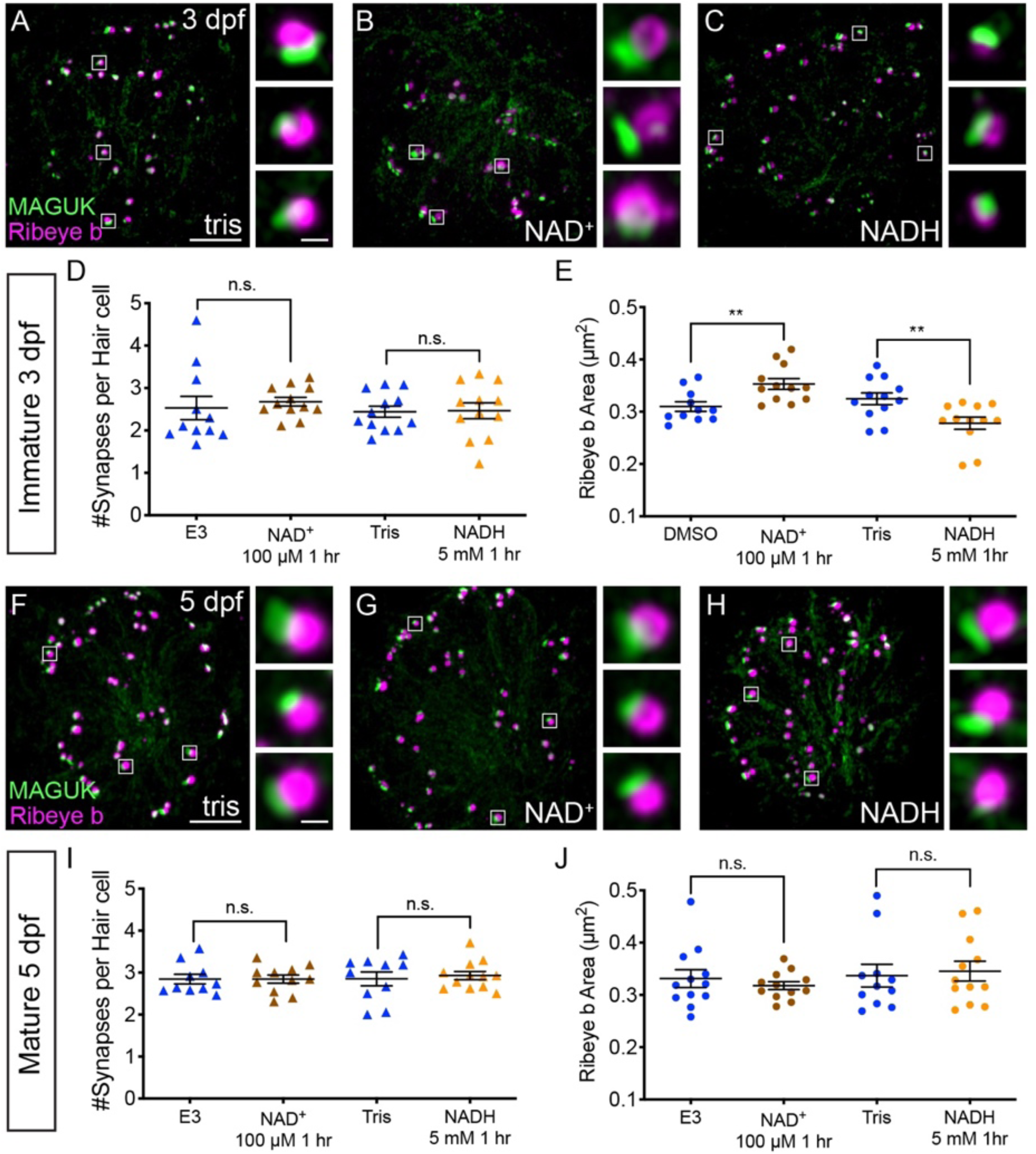
NAD^+^ and NADH directly influence ribbon formation. Representative images of immature (A-C, 3 dpf) and mature (G-H, 5 dpf) neuromasts immunostained with Ribeye b (magenta, ribbons) and MAGUK (green, postsynapses) after a 0.1% Tris-HCl (A, F), 100 μM NAD^+^ (B, G) or 5 mM NADH treatment (C, H). Insets show 3 example synapses (white squares). D-E and I-J, Scatterplots show synapse count (D, I) and ribbon area (E, J) in controls and treatments groups. N ≥ 10 neuromasts per treatment. Error bars in B-C represent SEM. A Welch’s unequal variance *t*-test was used for comparisons, \*\**p* < 0.01. Scale bar = 5 µm in A and F, 2 µm in insets.

## Discussion

In this study, we determined in a physiological setting how mito-Ca^2+^ influences hair-cell presynapse function and formation. In mature hair cells, evoked Ca_V_1.3-channel Ca^2+^ influx drives Ca^2+^ into mitochondria. Evoked mito-Ca^2+^ uptake is important to sustain presynaptic Ca^2+^ responses and maintain synapse integrity (Figure 8B). During development, spontaneous Ca_V_1.3 channel Ca^2+^ influx also drives Ca^2+^ into mitochondria. Elevated mito-Ca^2+^ levels rapidly lower the NAD^+^/NADH ratio and downregulate ribbon formation (Figure 8A). Furthermore, during development, NAD^+^ and NADH can directly increase and decrease ribbon formation respectively. Our study reveals an intriguing mechanism that couples presynaptic activity with mito-Ca^2+^ to regulate the function and formation of a presynaptic structure.

**Figure 8.**
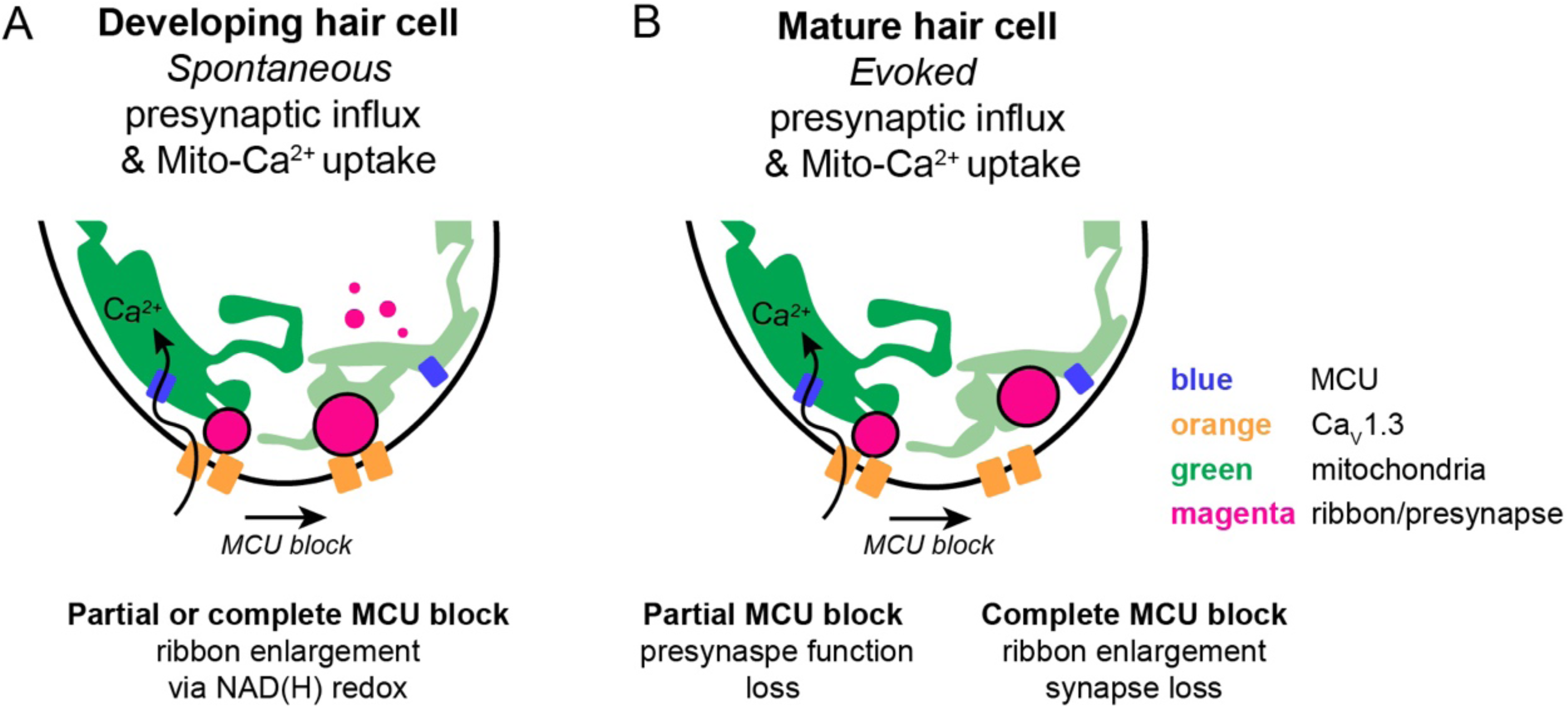
Schematic model of mito-Ca^2+^ in developing and mature hair cells. A, In developing hair cells, spontaneous presynaptic-Ca^2+^ influx is linked to mito-Ca^2+^ uptake. Together these Ca^2+^ signals function to regulate ribbon formation. When the Ca_V_1.3 or MCU channels are blocked, ribbon formation is increased leading to larger ribbons. These Ca^2+^ signals regulate ribbon formation via NAD(H) redox. B, In mature hair cells, evoked presynaptic-Ca^2+^ influx is linked to mito-Ca^2+^ uptake. When the MCU is blocked in mature hair cells there are synaptopathic consequences. Ribbons are enlarged and synapses are lost.

### Functional significance of ribbon size

Our work outlines how presynaptic activity controls the formation and ultimately the size of ribbons. When either presynaptic Ca^2+^ influx or mito-Ca^2+^ uptake was perturbed, ribbons were significantly larger (Figure 5A-C, E; <u>Sheets et al., 2012)</u>. But why regulate ribbon size?

Previous work has reported variations in ribbon size and shape among hair-cell types and species (Moser et al., 2006). Although ribbon morphology is predicted to impact synapse function, the functional consequence of presynapse structure on function has primarily been explored in the auditory inner hair cells of mice. In these auditory hair cells, studies have identified two distinct populations of ribbon synapses that spatially segregate within each cell (Kalluri and Monges-Hernandez, 2017; Liberman and Liberman, 2016; Liberman et al., 2011; Yin et al., 2014; Zhang et al., 2018a). Structurally, one population has smaller ribbons, while the other population has significantly larger ribbons. Functionally, compared to smaller ribbons, larger ribbons are associated with afferent fibers with less spontaneous activity and higher thresholds of activation <u>(Furman et al., 2013; Kalluri and Monges-Hernandez, 2017; Liberman et al., 2011, 2015, 1990; Merchan-Perez and Liberman, 1996; Song et al., 2016; Yin et al., 2014)</u>. Overall, the combined use of these two types of ribbon synapse is thought to increase the range of sensitivities for each individual auditory hair cell (Costalupes et al., 1984; Ohn et al., 2016). Interestingly, in mice these two populations of ribbons can be distinguished structurally just after the onset of hearing (Liberman and Liberman, 2016). This timing suggests that similar to our data (Figure 4-5), activity during development may help determine ribbon size.

Previous work in the zebrafish-lateral line has also examined how ribbon size impacts synapse function (Sheets et al., 2017). This work overexpressed Ribeye in zebrafish-hair cells to dramatically enlarge ribbons. Functionally, compared to controls, hair cells with enlarged ribbons were associated with afferent neurons with lower spontaneous activity (Sheets et al., 2017). Furthermore, the onset encoding, or the timing of the first afferent spike upon stimulation, was significantly delayed in hair cells with enlarged ribbons. Together, both studies in zebrafish and mouse indicate that ribbon size can impact the functional properties of the synapse. Based on these studies, we predict that the alterations to ribbon size we observed in our current study would impact functional properties of the synapse in a similar manner. For example, pharmacological treatments that enlarge ribbons (Figure 5: MCU channel block; Figure 7: exogenous NAD^+^) would also lower spontaneous spiking in afferents and delay onset encoding.

### Ribeye and CtBP localization at synapses

In this study, we found that NAD(H) redox state had a dramatic effect on ribbon formation. NAD^+^ promotes, while NADH reduces ribbon formation (Figure 7). The main component of ribbons is Ribeye. Ribeye has two domains, a unique A domain and a B domain that contains an NAD(H) binding domain (Schmitz et al., 2000a). *In vitro* work on isolated A and B domains has shown that both NAD^+^ and NADH can affect interactions between A and B domains as well as B-domain interactions (Magupalli et al., 2008). In the context of ribbons, the B domain has been shown to concentrate at the interface between the ribbon and the membrane opposing the postsynapse (Sheets et al., 2014). Therefore, promoting B domain homodimerization may act to seed larger ribbons at the presynapse. In this scenario, NAD^+^ and NADH could increase and decrease B domain homodimerization to impact ribbon formation. Because we also saw an increase in cytoplasmic Ribeye aggregates after MCU block (Figure 5F-G) it is alternatively possible that NAD^+^ and NADH could impact interactions between A and B domains to more broadly impact Ribeye interactions and accumulation.

Regardless of the exact mechanism, the effect of presynaptic activity and related changes in NAD(H) redox homeostasis may extend beyond the sensory ribbon synapse. Ribeye is a splice variant of the transcriptional co-repressor CtBP2 (Schmitz et al., 2000b). While the A domain is unique to Ribeye, the B domain is nearly identical to CtBP2 minus the nuclear localization sequence (NLS) (Hübler et al., 2012). In vertebrates, the CtBP family also includes CtBP1 (Chinnadurai, 2007). CtBP proteins are expressed in both hair cells and the nervous system, and there is evidence that both CtBP1 and CtBP2 may act as scaffolds at neuronal synapses (Hübler et al., 2012; tom Dieck et al., 2005). Interestingly, in cultured neurons, it has been shown that synaptic activity is associated with both an increase in CtBP1 localization at the presynapse, as well as a decrease in the NAD^+^/NADH ratio (Ivanova et al., 2015). In our *in vivo* study, we also found that the NAD^+^/NADH ratio was lower in developing hair cells with presynaptic activity (Figure 6H). But in contrast to *in vitro* work on CtBP1 in cultured neurons, we found that Ribeye localization to the presynapse and ribbon size was reduced when the NAD^+^/NADH ratio was lowered (Figure 7A-C). It is unclear why presynaptic activity regulates Ribeye localization differently from that of CtBP1. Ribeye and CtBP1 behavior may differ due to the divergent function of their N-terminal domains. Synaptic localization may also be influenced by external factors, such as the cell type in which the synapse operates, whether the study is performed *in vitro* or *in vivo*, as well as the maturity of the synapse. Overall, both studies demonstrate that CtBP1 and Ribeye localization to the presynapse can be influenced by synaptic activity and NAD(H) redox state.

### Role of evoked mito-Ca^2+^ uptake in mature hair cells

Studies in various neuronal subtypes have demonstrated that mitochondria play multiple roles to maintain neurotransmission including ATP production, Ca^2+^ buffering and signaling, and neurotransmitter synthesis. (reviewed in <u>Kann and Kovács, 2007; Vos et al.,</u> <u>2010)</u>. Our study found that in mature zebrafish-hair cells, even partially blocking evoked mito- Ca^2+^ uptake can impair presynaptic Ca^2+^ influx during sustained stimuli (Figure 2E-F). But how does mito-Ca^2+^ uptake impact presynaptic Ca^2+^ activity? Although mito-Ca^2+^ uptake could function to buffer cyto-Ca^2+^ to maintain presynaptic function, our current work indicates that blocking mito-Ca^2+^ uptake does not raise cytosolic Ca^2+^ levels (Figure 6A-C). Therefore mito-Ca^2+^ uptake may not be required to buffer or clear Ca^2+^ from the cytosol during steady-state. Alternatively, mito-Ca^2+^ uptake could buffer Ca^2+^ locally during presynaptic activity to prevent Ca^2+^-dependent inactivation of Ca_V_1.3 channels. In hair cells, Ca_V_1.3 channels exhibit reduced Ca^2+^ dependent inactivation (Koschak et al., 2001; Platzer et al., 2000; Song et al., 2003; Xu and Lipscombe, 2001). This reduction has been proposed to be important to transmit sustained sensory stimulation (Kollmar et al., 1997). Perhaps local removal of Ca^2+^ into the mitochondria during presynaptic activity is another mechanism in place to sustain neurotransmission. Alternatively, if mito-Ca^2+^ uptake does not buffer Ca^2+^, it could be critical to produce ATP for other cellular tasks to maintain neurotransmission. Additional work is necessary to understand how evoked mito-Ca^2+^ uptake functions to sustain presynaptic Ca^2+^ influx in mature-zebrafish hair cells.

In addition to innate cellular roles, in neurons and in hair cells, mito-Ca^2+^ loading is also associated with pathological processes such as reactive oxygen species (ROS) production, cell death and synapse loss (Cai and Tammineni, 2016; Court and Coleman, 2012; DiMauro and Schon, 2008; Esterberg et al., 2013, 2014; Sheng and Cai, 2012). Interestingly, recent work has demonstrated that noise-induced hearing loss is associated with measurable changes in ribbon morphology and synapse number (Jensen et al., 2015; Kujawa and Liberman, 2009; Liberman et al., 2015). Work studying this type of hearing loss has shown that auditory inner hair cells in the high frequency region of the mouse cochlea have enlarged ribbons immediately after noise, followed later by synapse loss (Liberman et al., 2015). This pathology is reminiscent of our 1-hr pharmacological treatments that completely block the MCU in mature zebrafish hair cells (Figure 3E-F). After this treatment, we observed a reduction in the number hair cells and synapses, and an increase in ribbon size. Surprisingly, these same treatments applied to developing hair cells increased ribbon size but did not reduce cell or synapse number (Figure 5D). Recent work has suggested that younger hair cells may be more resilient to ototoxins, perhaps because they have not yet accumulated an excess of mitochondria oxidation (Pickett et al., 2018). This could explain why complete MCU block is not pathological to developing hair cells. Overall these studies, along with our own data indicate that in mature hair cells, the mitochondria and the MCU may be associated with pathological processes associated with ototoxins and noise-exposure.

In further support of this idea, recent work in mice has investigated the role of the MCU in noise-related hearing loss (Wang et al., 2018). This work demonstrated that pharmacological block or a loss of function mutation in MCU protected against synapse loss in auditory inner hair cells after noise exposure. Although this result is counter to our observed results where complete MCU block reduces synapse number (Figure 3E), it highlights an association between mito-Ca^2+^, noise exposure and synapse integrity. It is possible that these differences can be explained by transitory versus chronic alterations in mito-Ca^2+^ homeostasis. These differences may be resolved by studying the hair cells in a zebrafish MCU knock out. In the future it will be interesting to examine both mito-Ca^2+^ uptake and ribbon morphology during other pathological conditions that enlarge ribbons such as noise exposure, ototoxicity and aging.

### Role of spontaneous mito-Ca^2+^ uptake in developing hair cells

Although mitochondria have been studied in the context of cellular function and cell death, relatively few studies have examined the role mitochondria play in development. We found that mitochondria spontaneously take up Ca^2+^ during hair-cell development (Figure 4B-C). Although studies in mammalian hair cells have demonstrated that there are spontaneous rises in presynaptic Ca^2+^ during development (Marcotti et al., 2003; Tritsch et al., 2010), these Ca^2+^ signals have not been reported in zebrafish hair cells. Our work highlights the mitochondria as a downstream signaling organelle that couples presynaptic-Ca^2+^ influx to ribbon formation (Figure 8A). In the future, zebrafish will be a useful model to further explore the origin and role of these spontaneous Ca^2+^ signals.

In our study, we also found that altering baseline mito-Ca^2+^ levels rapidly influenced the NAD^+^/NADH ratio and altered ribbon size in developing hair cells (Figure 5, 6, 7). However, in mature hair cells, while alterations to mito-Ca^2+^ levels increased ribbon size they did not influence NAD(H) redox (Figure 6I). One reason why NAD(H) redox does not change in mature hair cells is that ribbon enlargement could be occurring through a different mechanism. For example, ribbon enlargement could be a pathological byproduct of synapse loss (Figure 3E). In mature hair cells, after MCU block it is possible that individual ribbons are not enlarging, but instead ribbons are merging together as synapses are lost. In the future live imaging studies will help resolve whether there are different mechanisms underlying ribbon enlargement in mature and developing hair cells.

Overall this study has demonstrated the zebrafish-lateral line is a valuable system to study the interplay between the mitochondria, and synapse function, development and integrity. In the future it will be exciting to expand this research to explore how evoked and spontaneous mito-Ca^2+^ influx are impacted by pathological treatments such as age, noise and ototoxins.

## Method Details

### Zebrafish husbandry and genetics

Adult *Danio rerio* (zebrafish) were maintained under standard conditions. Larvae 2 to 6 days post-fertilization (dpf) were maintained in E3 embryo medium (in mM: 5 NaCl, 0.17 KCl, 0.33 CaCl_2_ and 0.33 MgSO_4_, buffered in HEPES pH 7.2) at 28°C. All husbandry and experiments were approved by the NIH Animal Care and Use program under protocol #1362-13. Transgenic zebrafish lines used in this study include: *Tg(myo6b:GCaMP6s-CAAX)^idc1^* (Jiang et al., 2017b), *Tg(myo6b:RGECO1)^vo10Tg^* (Maeda et al., 2014), Tg(myo6b:GCaMP3)^w78Tg^ (Esterberg et al., 2013), *Tg(myo6b:mitoGCaMP3)^w119Tg^* (Esterberg et al., 2014), and *Tg(myo6b:ribeye a-tagRFP)^idc11Tg^* (Sheets, 2017). Experiments were performed using Tübingen or TL wildtype strains.

### Cloning and Transgenic Fish Production

To create transgenic fish, plasmid construction was based on the tol2/Gateway zebrafish kit developed by the lab of Chi-Bin Chien at the University of Utah (Kwan et al., 2007). These methods were used to create *Tg(myo6b:mitoRGECO1)^idc12Tg^* and *Tg(myo6b:Rex-YFP)^idc13Tg^* transgenic lines. Gateway cloning was used to clone *Rex-YFP* (Bilan et al., 2014) and *mitoRGECO1* into the middle entry vector pDONR221. For mitochondrial matrix targeting, the sequence of cytochrome C oxidase subunit VIII (Rizzuto et al., 1989) was added to the N-terminus of RGECO1. Vectors p3E-polyA (Kwan et al., 2007) and pDestTol2CG2 (Kwan et al., 2007) were recombined with p5E *myosinVIb (myo6b)* (Kindt et al., 2012) and our engineered plasmids to create the following constructs: *myo6b:REX-YFP*, and *myo6b:mitoRGECO1.* To generate transgenic fish, DNA clones (25-50 ng/μl) were injected along with *tol2* transposase mRNA (25-50 ng/μl) into zebrafish embryos at the single-cell stage.

### Pharmacological treatment of larvae for immmunohistochemistry

For immunohistological studies, zebrafish larvae were exposed to compounds diluted in E3 with 0.1% DMSO (isradipine, Bay K8644, NAD^+^ (Sigma-Aldrich, St. Louis, MO), Ru360 (Millipore, Burlington, MA)) or Tris-HCl (NADH (Cayman Chemical, Ann Arbor, MI)) for 30 min or 1 hr at the concentrations indicated. E3 with 0.1% DMSO or Tris-HCl were used as control solutions. In solution at pH 7.0-7.3, NADH oxidizes into NAD^+^ by exposure to dissolved oxygen. To mitigate this, NADH was dissolved immediately before use, and was exchanged with a freshly dissolved NADH solution every half hour. Dosages of isradipine, Ru360, NAD^+^ and NADH did not confer excessive hair-cell death or synapse loss unless stated. After exposure to the compounds, larvae were quickly sedated on ice and transferred to fixative.

### In vivo imaging of baseline Ca^2+^ and NAD(H) redox

To prepare larvae for imaging, larvae were immobilized as previously described (Kindt et al, 2012). Briefly, larvae were anesthetized with tricaine (0.03%) and pinned to a chamber lined with Sylgard 184 Silicone Elastomer (Dow Corning, Midland, MI). Larvae were injected with 125 µM α-bungarotoxin (Tocris, Bristol, UK) into the pericardial cavity to paralyze. Tricaine was rinsed off the larvae with E3.

For baseline measurements of Rex-YFP and cytoRGECO1 fluorescence, larvae were imaged using an upright Nikon ECLIPSE Ni-E motorized microscope (Nikon Inc., Tokyo, Japan) in widefield mode with a Nikon 60x 1.0 NA CFI Fluor water-immersion objective, 480/30 nm excitation and 535/40 nm emission filter set or 520/35 nm excitation and 593/40 emission filter set, and an ORCA-D2 camera (Hamamatsu Photonics K.K., Hamamatsu City, Japan). Acquisitions were taken at 5 Hz, in 15 plane Z-stacks every 2 μm. For baseline measurements of MitoGCaMP3, larvae were imaged using a Bruker Swept-field confocal microscope (Bruker Inc., Billerica, MA), with a Nikon CFI Fluor 60x 1.0 NA water immersion objective. A Rolera EM-C2 CCD camera (QImaging, Surrey, Canada) was used to detect signals. Acquisitions were taken using a 70 µm slit at a frame rate of 10 Hz, in 26 plane Z-stacks every 1 μm. MitoGCaMP3 baseline intensity varied dramatically in controls between timepoints. To offset this variability, we acquired and averaged the intensity of 4 Z-stacks per time point. For all baseline measurements transgenic larvae were first imaged in E3 with 0.1% DMSO or 0.1% Tris-HCl as appropriate. Then larvae were exposed to pharmacological agents for 30 minutes and a second acquisition was taken. Any neuromasts with cell death after pharmacological treatment were excluded from our analyses.

### In vivo imaging of evoked Ca^2+^ signals

To measure evoked Ca^2+^ signals in hair-cells, larvae were prepared in a similar manner as described for baseline measurements. After α-bungarotoxin paralysis, larvae were immersed in neuronal buffer solution (in mM: 140 NaCl, 2 KCl, 2 CaCl_2_, 1 MgCl_2_ and 10 HEPES, pH 7.3). Evoked Ca^2+^ measurements were acquired using the Bruker Swept-field confocal system described above. To stimulate lateral-line hair cells, a fluid-jet was used as previously described to deliver a saturating stimulus (Lukasz and Kindt, 2018).

To measure presynaptic GCaMP6sCAAX signals at ribbons, images were acquired with 1 x 1 binning with a 35 µm slit at 50 Hz in a single plane containing presynaptic ribbons. Ribbons were marked in live hair cells using the *Tg(myo6b:ribeye a-tagRFP*)*^idc11Tg^* transgenic line (Figure S2). Ribbons were located relative to GCaMP6s signals by acquiring a Z-stack of 5 planes 1 μm. To correlate presynaptic GCaMP6sCAAX signals with mitoRGECO1 signals in hair cells, 2-color imaging was performed. Images were acquired in a single plane with 2 x 2 binning at 10 Hz. MitoGCaMP3 signals were acquired in Z-stacks of 5 planes 1 μm apart at 2 x 2 binning. High speed imaging along the Z-axis was accomplished by using a piezoelectric motor (PICMA P-882.11-888.11 series, Physik Instrumente GmbH, Karlsruhe, Germany) attached to the objective to allow rapid imaging at a 50 Hz frame rate yielding a 10 Hz volume rate. For pharmacological treatment, acquisitions were made prior to drug treatment and after a 20-min incubation in the pharmacological agent. Any neuromasts with cell death after pharmacological treatment were excluded from our analyses.

### In vivo imaging of spontaneous Ca^2+^ signals

To measure spontaneous Ca^2+^ signals in hair-cells, larvae were prepared in a similar manner as described for evoked Ca^2+^ measurements. Spontaneous Ca^2+^ measurements were acquired using the Bruker Swept-field confocal system described above. To measure spontaneous presynaptic GCaMP6sCAAX signals, images were acquired with 2 x 2 binning with a 70 µm slit at 0.33 Hz in a single plane for 900 s. For acquisition of two-color spontaneous presynaptic GCaMP6sCAAX and mitoRGECO1 signals images were acquired with 2 x 2 binning with a 70 µm slit at 0.2 Hz in a single plane for 900 s.

### Electron microscopy

Larvae were prepared for electron microscopy as described previously (Sheets, 2017). Transverse serial sections (∼60 nm thin sections) were used to section through neuromasts. Samples were imaged on a JEOL JEM-2100 electron microscope (JEOL Inc., Tokyo, Japan). The distance from the edge of a ribbon density to the edge of the nearest mitochondria was measured (n= 17 ribbons). In 74 % of ribbons, a mitochondrion could be clearly identified within 1 µm of a ribbon in a single section (17 out of 21 ribbons). All distances and perimeters were measured in FIJI (Schindelin et al., 2012).

### Immunofluorescence staining and Airyscan imaging

Whole larvae were fixed with 4% paraformaldehyde in PBS at 4°C for 3.5-4 hr as previously described (Zhang et al., 2018b). Fixative was washed out with 0.01% Tween in PBS (PBST) in 4 washes, 5 min each. Larvae were then washed for 5 min with H_2_O. The H_2_O was thoroughly removed and replaced with ice-cold acetone and placed at −20°C for 3 min for 3 dpf and 5 min for 5 dpf larvae, followed by a 5 min H_2_O wash. The larvae were then washed for 4 x 5 min in PBST, then incubated in block overnight at 4°C in blocking solution (2% goat serum, 1 % bovine serum albumin, 2% fish skin gelatin in PBST). Primary and secondary antibodies were diluted in blocking solution. Primary antibodies and their respective dilution are: Ribbon label: Mouse anti-Ribeye b IgG2a, 1:10,000 (Sheets et al., 2011b); PSD label: Mouse anti-pan-MAGUK IgG1 #75-029, 1:500 (UC Davis/NIH NeuroMab Facility, Davis, CA); Hair cell label: Rabbit anti-Myosin VIIa, 1:1000 (Proteus BioSciences Inc., Ramona, CA). Larvae were incubated in primary antibody solution for 2 hr at room temperature. After 4 x 5 min washes in PBST to remove the primary antibodies, diluted secondary antibodies were added in and samples were incubated for 2 hr at room temperature. Secondary antibodies and their respective dilution are: goat anti-mouse IgG2a, Alexa Fluor 488, 1:1000; goat anti-rabbit IgG (H+L) Alexa Fluor 568, 1:1000; goat anti-mouse IgG1 Alexa Fluor 647, 1:1000 (Thermo Fisher Scientific, Waltham, MA). Secondary antibody was washed out with PBST for 3 x 5 min, followed by a 5 min wash with H_2_O. Larvae were mounted on glass slides with Prolong Gold Antifade Reagent (Invitrogen, Carlsbad, CA) using No. 1.5 coverslips.

Prior to Airyscan imaging, live samples were immobilized in 2 % low-melt agarose in tricaine (0.03%) in cover-glass bottomed dishes. Live and fixed samples were imaged on an inverted Zeiss LSM 780 laser-scanning confocal microscope with an Airyscan attachment (Carl Zeiss AG, Oberkochen, Germany) using an 63x 1.4 NA oil objective lens. The median (± median absolute deviation) lateral and axial resolution of the system was measured at 198 ± 7.5 nm and 913 ± 50 nm (full-width at half-maximum), respectively. The acquisition parameters were adjusted using the control sample such that pixels for each channel reach at least 1/10 of the dynamic range. The Airyscan Z-stacks were processed with Zeiss Zen Black software v2.1 using 3D filter setting of 7.0. Experiments were imaged with the same acquisition settings to maintain consistency between comparisons.

## Quantification and Statistical Analysis

### Analysis of Ca^2+^and NAD(H) signals, processing, and quantification

To quantify changes in baseline Ca^2+^ and NAD(H) homeostasis, images were processed in FIJI. For our measurements we quantified the fluorescence in the basal-most 8 μm (4 planes) to avoid overlap between cells. The basal planes were max Z-projected, and a 24.0μm (Rex-YFP and RGECO1) or 26.8 μm (MitoGCaMP3) circular region of interest (ROI) was drawn over the neuromast to make an intensity measurement. To correct for photobleaching, a set of mock-treated control neuromasts were imaged during every trial. These mock treatments were used to normalize the post-treatment intensity values.

To quantify the magnitude of evoked changes in Ca^2+^, fluorescent images were processed in FIJI. Images in each time series were aligned using Stackreg (Thevenaz et al., 1998). For evoked MitoRGECO1, MitoGCaMP3, CytoGCaMP3 and two-color GCaMP6sCAAX and MitoRGECO1 signals, Z-stack were max z-projected, and a 5 μm diameter circular ROI was drawn over each hair cell. For ribbon-localized measurements, GCaMP6sCAAX signals were measured within a 1.34 μm round ROIs at individual ribbons, and intensity of multiple ROI within a cell were averaged. Cells with presynaptic Ca^2+^ activity is defined by max ΔF/F of > 0.05 for MitoRGECO1 and MitoGCaMP3, and max ΔF/F > 0.25 for GCaMP6sCAAX for a 2-s stimulation.

To quantify the average magnitude and frequency of spontaneous Ca^2+^ changes in GCaMP6sCAAX signals, images were processed in Matlab R2014b (Mathworks, Natick, MA) and FIJI. First, images in each time series were aligned using Stackreg (Thevenaz et al., 1998). To measure the average magnitude during the 900 s GCaMP6sCAAX image acquisition, a 5 μm diameter circular ROI was drawn over each hair cell and a raw intensity value was obtained from each time point. Then the raw traces were bleach corrected. Next, the corrected intensity values were normalized as ΔF/F_o_. F_o_ is defined as the bottom 15^th^ percentile of fluorescence values (Babola et al., 2018). Then, values of ΔF/F_0_ of less than 10 % were removed. These values were considered to be noise and our threshold value for a true signal. A 10 % threshold was determined by imaging spontaneous GCaMP6CAAX signals in the presence of isradipine where no signals were observed (Figure S4). The averaged magnitude of spontaneous activity per cell was obtained by dividing the integral/sum of GCaMP6sCAAX signals (ΔF/F_o_ > 10%) during the whole recording period by 300 (300 frames in 900 s). The frequency of GCaMP6sCAAX signals was defined as the average number of peaks per second during the whole recording period.

### Image processing and quantification of synapse morphology

To quantify synapse morphology and pairing, images were first processed in ImageJ (NIH, Bethesda, MD), and then synapses were paired using Python (Python Software Foundation, Wilmington, DE) in the Spyder Scientific Environment (MIT, Cambridge, MA). In ImageJ, each Airyscan Z-stack was background subtracted using rolling-ball subtraction. Z-stacks containing the MAGUK channel were further bandpass filtered to remove details smaller than 6 px and larger than 20 px. A duplicate of the Z-stack was normalized for intensity. This duplicated Z-stack was used to identify individual ribbon and MAGUK using the Simple 3D Segmentation of ImageJ 3D Suite (Ollion et al., 2013). Local intensity maxima, identified with 3D Fast Filter, and 3D watershed were used to separate close-by structures. The centroid for each identified ribbon and MAGUK was obtained using 3D Manager and were used to identify complete synapses. The max Z-projection of the segmented Z-stack was used to generate a list of 2D objects as individual ROIs corresponding to each punctum. This step also included a minimum size filter, Ribeye: 0.08 μm^2^, MAGUK 0.04 μm^2^. For quantification of extrasynaptic Ribeye b puncta, the minimum size filter was not applied. The 2D puncta ROI were applied over the max Z-projection of the original Z-stack processed only with background subtraction. This step measures the intensity of the antibody label. Centroid and intensity information were exported as a CSV spreadsheet (macro is available upon request).

In Python, the 3D centroid coordinates for each ribbon punctum was measured against the coordinates of every post-synaptic MAGUK punctum to find the MAGUK punctum within a threshold distance. This threshold was calculated by taking the 2D area of the Ribeye and MAGUK punctum measured in the max Z-projection to calculate an approximate radius by dividing by π and taking the square root. The two radii were then summed to get the threshold. Puncta that were not paired were excluded from later statistical analyses of synaptic ribbon and postsynaptic MAGUK puncta. Hair cell and synapse count were confirmed manually. Hair cell counts were performed with myosin VIIa antibody label in treatments where synapse or cell numbers were reduced.

### Statistics

Statistical analyses and data plots were performed with Prism 8 (Graphpad, San Diego, CA). Values in the text and data with error bars on graphs and in text are expressed as mean ± SEM unless indicated otherwise. All experiments were performed on a minimum of 2 animals, 6 neuromasts (posterior lateral-line neuromasts L1-L4), on 2 independent days. For 3 and 5 dpf larvae each neuromast represents analysis from 8-12 hair cells; 24-36 synapses and 14-18 hair cells; 42-54 synapses respectively. All replicates are biological. Based on the variance and effect sizes reported previously and measured in this study, these numbers were adequate to provide statistical power to avoid both Type I and Type II error (Sheets et al., 2012; Zhang et al., 2018b). No animals or samples were excluded from our analyses unless control experiments failed–in these cases all samples were excluded. No randomization or blinding was used for our animal studies. Where appropriate, datasets were confirmed for normality using a D’Agostino-Pearson normality test and for equal variances using a F test to compare variances. Statistical significance between two conditions was determined by either unpaired Welch’s unequal variance *t*-test, a Mann-Whitney U test or a Wilcoxon matched-pairs signed-rank test as appropriate. For comparison of multiple conditions, a Brown-Forsythe or a Welch ANOVA with Games-Howell post hoc were used.

## Acknowledgements

This work was supported by National Institute on Deafness and Other Communication Disorders Intramural Research Program Grant 1ZIADC000085-01 to K.S.K. and ZICDC000081 to R.S.P. and Y.-X.W. We would like to thank Katie Drerup, Paul Fuchs and Doris Wu for their support and thoughtful comments on the manuscript.

## Author contributions

Conceptualization, Methodology, Writing, H.C.W., K.S.K., Formal Analysis, H.C.W., Investigation, H.C.W., Q.X.Z., A.J.B., R.S.P., Y.-X.W., and Supervision, K.S.K.

---

**Movie S1.** Airyscan image of MitoGCaMP3 and Rib a-tagRFP at the base of a single live hair cell.

**Movie S2.** Spontaneous ΔF GCaMP6sCAAX (left) and ΔF MitoRGECO1 (right) signals acquired at 3 dpf, 25-s per frame.

**Figure S1.**
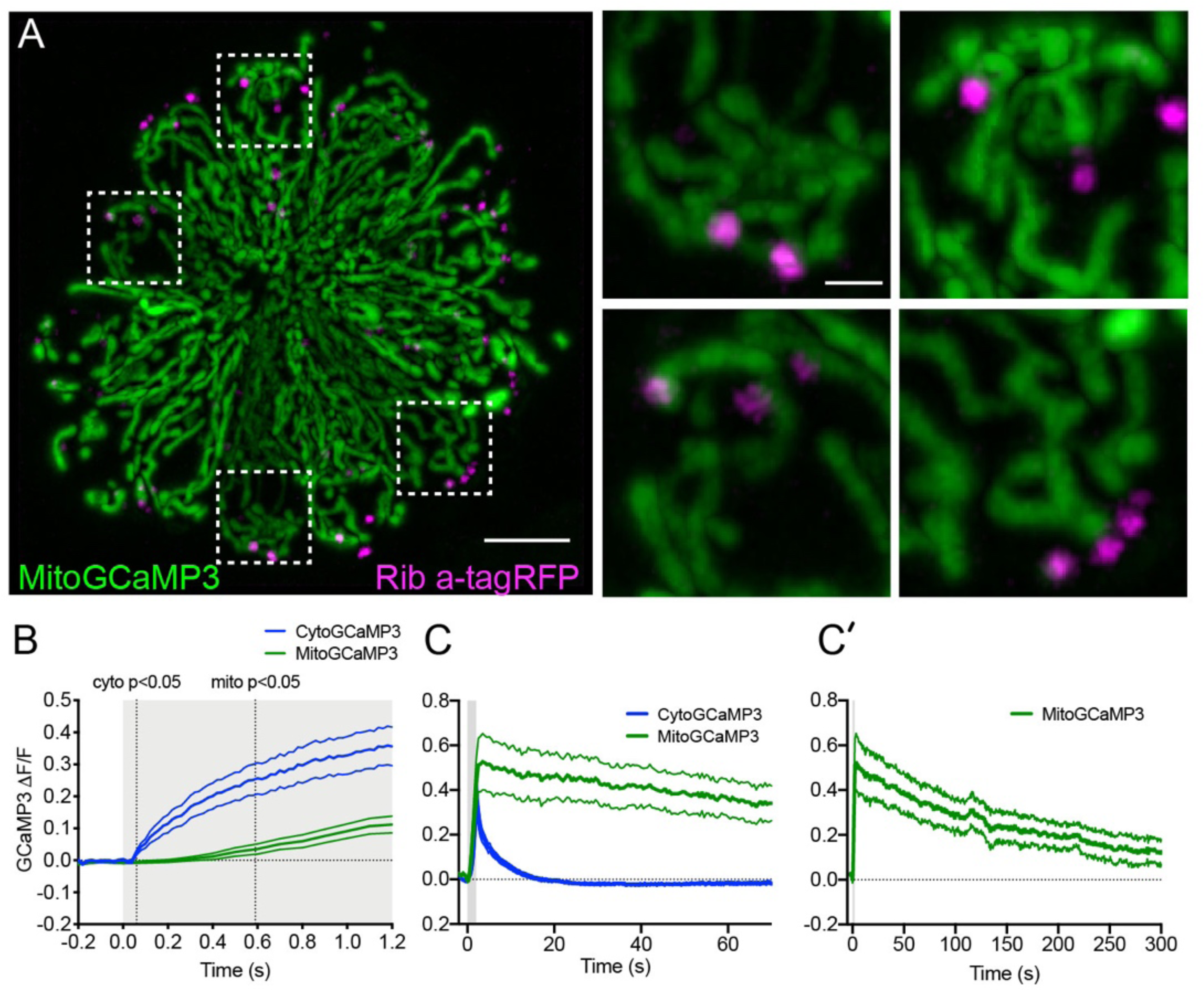
The time course of mechanically-evoked mito-Ca^2+^ and cyto-Ca^2+^ signals are distinct. A, Airyscan confocal image of a live, neuromast expressing MitoGCaMP3 (mitochondria) and Ribeye a-tagRFP (ribbons) at 6 dpf. Insets show the base of 4 individual hair cells from the neuromast in *A* (dashed white boxes). B, Average cyto-(blue) and mito-Ca^2+^ (green) signals during the onset of a 2-s stimulus. Mito-Ca^2+^ signals rise with a delay compared to cyto-Ca^2+^ signals (3-6 dpf, n ≥ 18 cells). C-C’, Average cyto- and mito-Ca^2+^ signals during and after a 2-s stimulation shows that cyto-Ca^2+^ signals return to baseline shortly after stimulation (C), while mito-Ca^2+^ remains elevated up to 5 min after stimulation (C-C’) (3 dpf, n ≥ 7 cells). Error in panel B-C’ represent SEM. Scale bar = 5 µm in A and 2 µm in inset.

**Figure S2.**
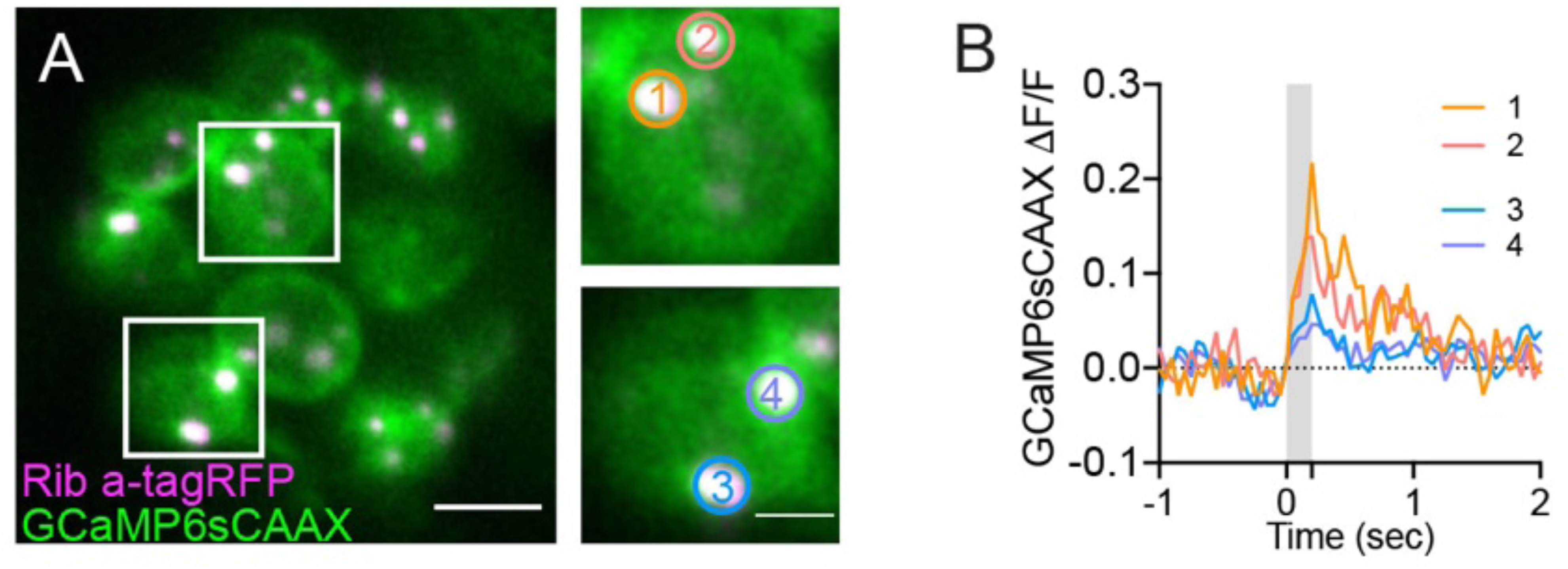
Presynaptic Ca^2+^ signals at the ribbon synapse. A, Live image of a neuromast viewed top-down, expressing the presynaptic-Ca^2+^ sensor GCaMP6sCAAX (green) and ribbon label Ribeye a-tagRFP (magenta) at 3 dpf. Example cells show evoked synaptic-Ca^2+^ signals during a 0.2-s stimulation (white boxes, duplicated on right). Circles 1-4 (1.3 μm diameter) denote regions used to generate the temporal traces of presynaptic-Ca^2+^ signals in *B*. Scale bar = 5 µm in A and 2 µm in insets.

**Figure S3.**
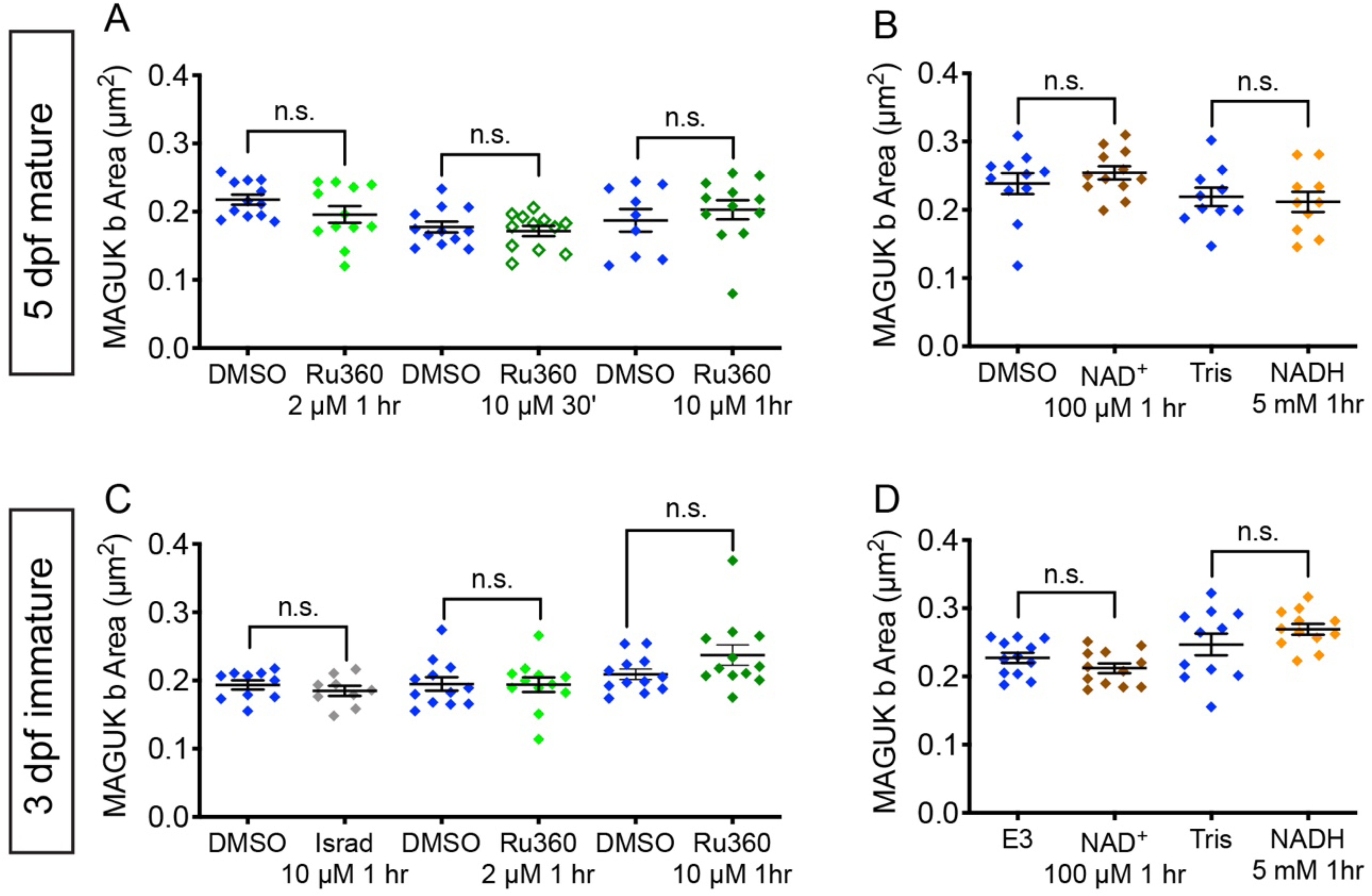
NAD^+^, NADH and Ru360 treatments do not impact postsynapse size. (A-D). Quantification of postsynapse size assayed by MAGUK immunolabel in mature (A-B) and developing neuromasts (C-D). Treatments with E3, 0.1% DMSO, 0.1% Tris-HCl, 100 μM NAD^+^, 5 mM NADH treatment, 2 µM Ru360, 10 µM Ru360 do not significantly alter postsynapse size compared to controls. (C, H). N ≥ 9 neuromasts per treatment. Error bars in B-C represent SEM. A Welch’s unequal variance *t*-test was used for comparisons.

**Figure S4.**
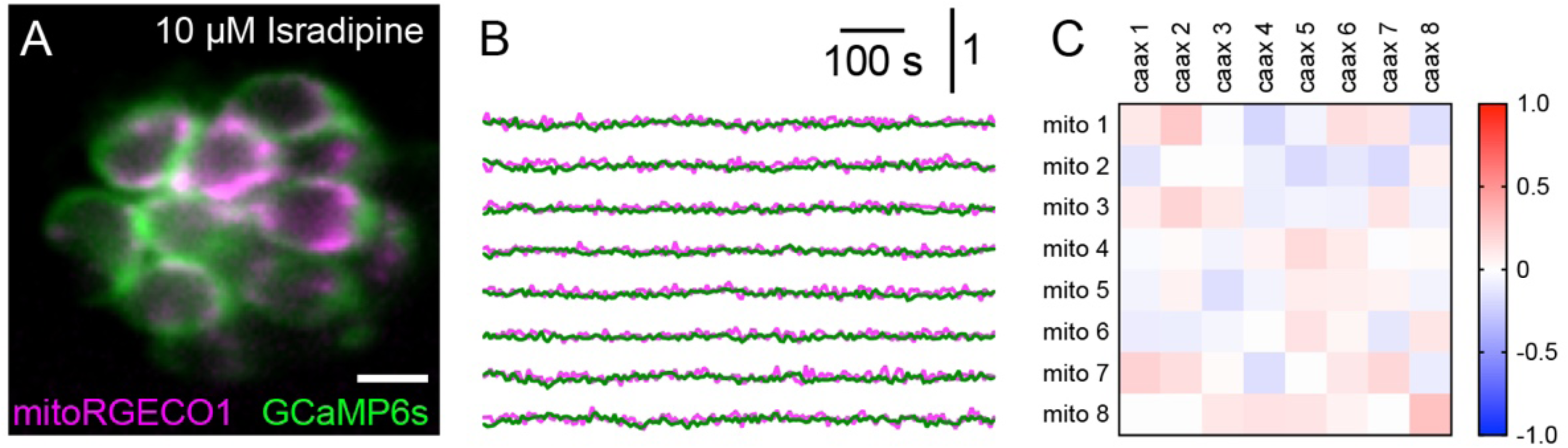
Spontaneous presynaptic and mito-Ca^2+^ signals are abolished by Ca_V_1.3 channel antagonist isradipine. A, A live Image of a neuromast viewed top-down, expressing the presynaptic-Ca^2+^ sensor GCaMP6sCAAX (green) and mito-Ca^2+^ sensor MitoRGECO1 (magenta) at 6 dpf. B, Representative GCaMP6sCAAX (green) and MitoRGECO1 (magenta) traces during a 900-s continuous image acquisition in the absence of stimuli and 10 µM isradipine. C, There is no correlation between GCaMP6sCAAX and MitoRGECO1 signals within each cell in the presence of isradipine.

## References

Babola, T.A., Li, S., Gribizis, A., Lee, B.J., Issa, J.B., Wang, H.C., Crair, M.C., and Bergles, D.E. (2018). Homeostatic Control of Spontaneous Activity in the Developing Auditory System. Neuron 99, 511–524.e5.

Becker, L., Schnee, M.E., Niwa, M., Sun, W., Maxeiner, S., Talaei, S., Kachar, B., Rutherford, M.A., and Ricci, A.J. (2018). The presynaptic ribbon maintains vesicle populations at the hair cell afferent fiber synapse. ELife Sciences 7, e30241.

Billups, B., and Forsythe, I.D. (2002). Presynaptic mitochondrial calcium sequestration influences transmission at mammalian central synapses. The Journal of Neuroscience : The Official Journal of the Society for Neuroscience 22, 5840–5847.

Böttger, E.C., and Schacht, J. (2013). The mitochondrion: a perpetrator of acquired hearing loss. Hear. Res. 303, 12–19.

Brandt, A., Striessnig, J., and Moser, T. (2003). CaV1.3 channels are essential for development and presynaptic activity of cochlear inner hair cells. J. Neurosci. 23, 10832–10840.

Cai, Q., and Tammineni, P. (2016). Alterations in Mitochondrial Quality Control in Alzheimer’s Disease. Front. Cell. Neurosci. 10.

Carafoli, E. (2011). The plasma membrane calcium pump in the hearing process: physiology and pathology. Sci China Life Sci 54, 686–690.

Castellano-Muñoz, M., and Ricci, A.J. (2014). Role of intracellular calcium stores in hair-cell ribbon synapse. Front Cell Neurosci 8.

Ceriani, F., Hendry, A., Jeng, J.-Y., Johnson, S.L., Stephani, F., Olt, J., Holley, M.C., Mammano, F., Engel, J., Kros, C.J., et al. (2019). Coordinated calcium signalling in cochlear sensory and non-sensory cells refines afferent innervation of outer hair cells. The EMBO Journal e99839.

Chinnadurai, G. (2007). Transcriptional regulation by C-terminal binding proteins. The International Journal of Biochemistry & Cell Biology 39, 1593–1607.

Chouhan, A.K., Zhang, J., Zinsmaier, K.E., and Macleod, G.T. (2010). Presynaptic mitochondria in functionally different motor neurons exhibit similar affinities for Ca2+ but exert little influence as Ca2+ buffers at nerve firing rates in situ. The Journal of Neuroscience : The Official Journal of the Society for Neuroscience 30, 1869–1881.

Costalupes, J.A., Young, E.D., and Gibson, D.J. (1984). Effects of continuous noise backgrounds on rate response of auditory nerve fibers in cat. Journal of Neurophysiology 51, 1326–1344.

Court, F.A., and Coleman, M.P. (2012). Mitochondria as a central sensor for axonal degenerative stimuli. Trends Neurosci. 35, 364–372.

Devine, M.J., and Kittler, J.T. (2018). Mitochondria at the neuronal presynapse in health and disease. Nature Reviews Neuroscience 19, 63–80.

DiMauro, S., and Schon, E.A. (2008). Mitochondrial Disorders in the Nervous System. Annual Review of Neuroscience 31, 91–123.

Eckrich, T., Blum, K., Milenkovic, I., and Engel, J. (2018). Fast Ca2+ Transients of Inner Hair Cells Arise Coupled and Uncoupled to Ca2+ Waves of Inner Supporting Cells in the Developing Mouse Cochlea. Front. Mol. Neurosci. 11.

Esterberg, R., Hailey, D.W., Coffin, A.B., Raible, D.W., and Rubel, E.W. (2013). Disruption of Intracellular Calcium Regulation Is Integral to Aminoglycoside-Induced Hair Cell Death. Journal of Neuroscience 33, 7513–7525.

Esterberg, R., Hailey, D.W., Rubel, E.W., and Raible, D.W. (2014). ER–Mitochondrial Calcium Flow Underlies Vulnerability of Mechanosensory Hair Cells to Damage. J Neurosci 34, 9703– 9719.

Fischel-Ghodsian, N., Kopke, R.D., and Ge, X. (2004). Mitochondrial dysfunction in hearing loss. Mitochondrion 4, 675–694.

Fjeld, C.C., Birdsong, W.T., and Goodman, R.H. (2003). Differential binding of NAD+ and NADH allows the transcriptional corepressor carboxyl-terminal binding protein to serve as a metabolic sensor. Proc. Natl. Acad. Sci. U.S.A. 100, 9202–9207.

Flippo, K.H., and Strack, S. (2017). Mitochondrial dynamics in neuronal injury, development and plasticity. Journal of Cell Science 130, 671–681.

Furman, A.C., Kujawa, S.G., and Liberman, M.C. (2013). Noise-induced cochlear neuropathy is selective for fibers with low spontaneous rates. J. Neurophysiol. 110, 577–586.

Hübler, D., Rankovic, M., Richter, K., Lazarevic, V., Altrock, W.D., Fischer, K.-D., Gundelfinger, E.D., and Fejtova, A. (2012). Differential spatial expression and subcellular localization of CtBP family members in rodent brain. PloS One 7, e39710.

Ivanova, D., Dirks, A., Montenegro-Venegas, C., Schöne, C., Altrock, W.D., Marini, C., Frischknecht, R., Schanze, D., Zenker, M., Gundelfinger, E.D., et al. (2015). Synaptic activity controls localization and function of CtBP1 via binding to Bassoon and Piccolo. EMBO J 34, 1056–1077.

Jean, P., Morena, D.L. de la, Michanski, S., Tobón, L.M.J., Chakrabarti, R., Picher, M.M., Neef, J., Jung, S., Gültas, M., Maxeiner, S., et al. (2018). The synaptic ribbon is critical for sound encoding at high rates and with temporal precision. ELife Sciences 7, e29275.

Jensen, J.B., Lysaght, A.C., Liberman, M.C., Qvortrup, K., and Stankovic, K.M. (2015). Immediate and delayed cochlear neuropathy after noise exposure in pubescent mice. PloS One 10, e0125160.

Jensen-Smith, H.C., Hallworth, R., and Nichols, M.G. (2012). Gentamicin rapidly inhibits mitochondrial metabolism in high-frequency cochlear outer hair cells. PLoS ONE 7, e38471.

Jiang, T., Kindt, K., and Wu, D.K. (2017a). Transcription factor Emx2 controls stereociliary bundle orientation of sensory hair cells. ELife 6.

Jiang, T., Kindt, K., and Wu, D.K. (2017b). Transcription factor Emx2 controls stereociliary bundle orientation of sensory hair cells. ELife 6.

Johnson, S.L., Safieddine, S., Mustapha, M., and Marcotti, W. (2019). Hair Cell Afferent Synapses: Function and Dysfunction. Cold Spring Harb Perspect Med.

Kalluri, R., and Monges-Hernandez, M. (2017). Spatial Gradients in the Size of Inner Hair Cell Ribbons Emerge Before the Onset of Hearing in Rats. Journal of the Association for Research in Otolaryngology 18, 399–413.

Kann, O., and Kovács, R. (2007). Mitochondria and neuronal activity. Am. J. Physiol., Cell Physiol. 292, C641–57.

Kennedy, H.J. (2002). Intracellular calcium regulation in inner hair cells from neonatal mice. Cell Calcium 31, 127–136.

Kindt, K.S., Finch, G., and Nicolson, T. (2012). Kinocilia mediate mechanosensitivity in developing zebrafish hair cells. Developmental Cell 23, 329–341.

Kokotas, H., Petersen, M.B., and Willems, P.J. (2007). Mitochondrial deafness. Clin. Genet. 71, 379–391.

Kollmar, R., Fak, J., Montgomery, L.G., and Hudspeth, A.J. (1997). Hair cell-specific splicing of mRNA for the alpha1D subunit of voltage-gated Ca2+ channels in the chicken’s cochlea. Proc. Natl. Acad. Sci. U.S.A. 94, 14889–14893.

Koschak, A., Reimer, D., Huber, I., Grabner, M., Glossmann, H., Engel, J., and Striessnig, J. (2001). alpha 1D (Cav1.3) subunits can form l-type Ca2+ channels activating at negative voltages. J. Biol. Chem. 276, 22100–22106.

Kujawa, S.G., and Liberman, M.C. (2009). Adding insult to injury: cochlear nerve degeneration after “temporary” noise-induced hearing loss. The Journal of Neuroscience : The Official Journal of the Society for Neuroscience 29, 14077–14085.

Kwan, K.M., Fujimoto, E., Grabher, C., Mangum, B.D., Hardy, M.E., Campbell, D.S., Parant, J.M., Yost, H.J., Kanki, J.P., and Chien, C.-B. (2007). The Tol2kit: a multisite gateway-based construction kit for Tol2 transposon transgenesis constructs. Dev. Dyn. 236, 3088–3099.

Kwon, S.-K., Sando, R., Lewis, T.L., Hirabayashi, Y., Maximov, A., and Polleux, F. (2016). LKB1 Regulates Mitochondria-Dependent Presynaptic Calcium Clearance and Neurotransmitter Release Properties at Excitatory Synapses along Cortical Axons. PLOS Biology 14, e1002516.

Lagnado, L., and Schmitz, F. (2015). Ribbon Synapses and Visual Processing in the Retina. Annual Review of Vision Science 1, 235–262.

Lepeta, K., Lourenco, M.V., Schweitzer, B.C., Martino Adami, P.V., Banerjee, P., Catuara-Solarz, S., de La Fuente Revenga, M., Guillem, A.M., Haidar, M., Ijomone, O.M., et al. (2016). Synaptopathies: synaptic dysfunction in neurological disorders – A review from students to students. J Neurochem 138, 785–805.

Levy, M., Faas, G.C., Saggau, P., Craigen, W.J., and Sweatt, J.D. (2003). Mitochondrial Regulation of Synaptic Plasticity in the Hippocampus. Journal of Biological Chemistry 278, 17727–17734.

Liberman, L.D., and Liberman, M.C. (2016). Postnatal maturation of auditory-nerve heterogeneity, as seen in spatial gradients of synapse morphology in the inner hair cell area. Hearing Research 339, 12–22.

Liberman, L.D., Wang, H., and Liberman, M.C. (2011). Opposing Gradients of Ribbon Size and AMPA Receptor Expression Underlie Sensitivity Differences among Cochlear-Nerve/Hair-Cell Synapses. J Neurosci 31, 801–808.

Liberman, L.D., Liberman, M.C., and Liberman, M.C. (2015). Dynamics of cochlear synaptopathy after acoustic overexposure. Journal of the Association for Research in Otolaryngology 16, 205– 219.

Liberman, M.C., Dodds, L.W., and Pierce, S. (1990). Afferent and efferent innervation of the cat cochlea: quantitative analysis with light and electron microscopy. The Journal of Comparative Neurology 301, 443–60.

Lioudyno, M., Hiel, H., Kong, J.-H., Katz, E., Waldman, E., Parameshwaran-Iyer, S., Glowatzki, E., and Fuchs, P.A. (2004). A “synaptoplasmic cistern” mediates rapid inhibition of cochlear hair cells. J. Neurosci. 24, 11160–11164.

Llorente-Folch, I., Rueda, C.B., Pardo, B., Szabadkai, G., Duchen, M.R., and Satrustegui, J. (2015). The regulation of neuronal mitochondrial metabolism by calcium. J Physiol 593, 3447–3462.

Lukasz, D., and Kindt, K.S. (2018). In Vivo Calcium Imaging of Lateral-line Hair Cells in Larval Zebrafish. J Vis Exp.

Maeda, R., Kindt, K.S., Mo, W., Morgan, C.P., Erickson, T., Zhao, H., Clemens-Grisham, R., Barr-Gillespie, P.G., and Nicolson, T. (2014). Tip-link protein protocadherin 15 interacts with transmembrane channel-like proteins TMC1 and TMC2. Proceedings of the National Academy of Sciences 111, 12907–12912.

Magupalli, V.G., Schwarz, K., Alpadi, K., Natarajan, S., Seigel, G.M., and Schmitz, F. (2008). Multiple RIBEYE-RIBEYE interactions create a dynamic scaffold for the formation of synaptic ribbons. The Journal of Neuroscience : The Official Journal of the Society for Neuroscience 28, 7954–7967.

Marcotti, W., Johnson, S.L., Rusch, A., and Kros, C.J. (2003). Sodium and calcium currents shape action potentials in immature mouse inner hair cells. J. Physiol. (Lond.) 552, 743–761.

Matlib, M.A., Zhou, Z., Knight, S., Ahmed, S., Choi, K.M., Krause-Bauer, J., Phillips, R., Altschuld, R., Katsube, Y., Sperelakis, N., et al. (1998). Oxygen-bridged dinuclear ruthenium amine complex specifically inhibits Ca2+ uptake into mitochondria in vitro and in situ in single cardiac myocytes. Journal of Biological Chemistry 273, 10223–10231.

Matthews, G., and Fuchs, P. (2010). The diverse roles of ribbon synapses in sensory neurotransmission. Nat. Rev. Neurosci. 11, 812–822.

McHenry, M.J., Feitl, K.E., Strother, J.A., and Van Trump, W.J. (2009). Larval zebrafish rapidly sense the water flow of a predator’s strike. Biology Letters 5, 477–479.

Merchan-Perez, A., and Liberman, M.C. (1996). Ultrastructural differences among afferent synapses on cochlear hair cells: correlations with spontaneous discharge rate. J. Comp. Neurol. 371, 208–221.

Metcalfe, W.K. (1985). Sensory neuron growth cones comigrate with posterior lateral line primordial cells in zebrafish. The Journal of Comparative Neurology 238, 218–224.

Moser, T., Brandt, A., and Lysakowski, A. (2006). Hair cell ribbon synapses. Cell Tissue Res 326, 347–359.

Mulkey, R.M., and Malenka, R.C. (1992). Mechanisms underlying induction of homosynaptic long-term depression in area CA1 of the hippocampus. Neuron 9, 967–975.

Murakami, S.L., Cunningham, L.L., Werner, L.A., Bauer, E., Pujol, R., Raible, D.W., and Rubel, E.W. (2003). Developmental differences in susceptibility to neomycin-induced hair cell death in the lateral line neuromasts of zebrafish (Danio rerio). Hearing Research 186, 47–56.

Ohn, T.-L., Rutherford, M.A., Jing, Z., Jung, S., Duque-Afonso, C.J., Hoch, G., Picher, M.M., Scharinger, A., Strenzke, N., and Moser, T. (2016). Hair cells use active zones with different voltage dependence of Ca2+ influx to decompose sounds into complementary neural codes. Proceedings of the National Academy of Sciences 113, E4716–E4725.

Ollion, J., Cochennec, J., Loll, F., Escudé, C., and Boudier, T. (2013). TANGO: a generic tool for high-throughput 3D image analysis for studying nuclear organization. Bioinformatics 29, 1840– 1841.

Pickett, S.B., Thomas, E.D., Sebe, J.Y., Linbo, T., Esterberg, R., Hailey, D.W., and Raible, D.W. (2018). Cumulative mitochondrial activity correlates with ototoxin susceptibility in zebrafish mechanosensory hair cells. Elife 7.

Platzer, J., Engel, J., Schrott-Fischer, A., Stephan, K., Bova, S., Chen, H., Zheng, H., and Striessnig, J. (2000). Congenital deafness and sinoatrial node dysfunction in mice lacking class D L-type Ca2+ channels. Cell 102, 89–97.

Rizzuto, R., Nakase, H., Darras, B., Francke, U., Fabrizi, G.M., Mengel, T., Walsh, F., Kadenbach, B., DiMauro, S., and Schon, E.A. (1989). A gene specifying subunit VIII of human cytochrome c oxidase is localized to chromosome 11 and is expressed in both muscle and non-muscle tissues. J. Biol. Chem. 264, 10595–10600.

Safieddine, S., El-Amraoui, A., and Petit, C. (2012). The Auditory Hair Cell Ribbon Synapse: From Assembly to Function. Annual Review of Neuroscience 35, 509–528.

Santos, F., MacDonald, G., Rubel, E.W., and Raible, D.W. (2006). Lateral line hair cell maturation is a determinant of aminoglycoside susceptibility in zebrafish (Danio rerio). Hearing Research 213, 25–33.

Schindelin, J., Arganda-Carreras, I., Frise, E., Kaynig, V., Longair, M., Pietzsch, T., Preibisch, S., Rueden, C., Saalfeld, S., Schmid, B., et al. (2012). Fiji: an open-source platform for biological-image analysis. Nat. Methods 9, 676–682.

Schmitz, F., Königstorfer, A., and Südhof, T.C. (2000a). RIBEYE, a component of synaptic ribbons: a protein’s journey through evolution provides insight into synaptic ribbon function. Neuron 28, 857–872.

Schmitz, F., Königstorfer, A., and Südhof, T.C. (2000b). RIBEYE, a component of synaptic ribbons: a protein’s journey through evolution provides insight into synaptic ribbon function. Neuron 28, 857–872.

Schnee, M.E., and Ricci, A.J. (2003). Biophysical and pharmacological characterization of voltage-gated calcium currents in turtle auditory hair cells. J. Physiol. (Lond.) 549, 697–717.

Sheets, L. (2017). Excessive activation of ionotropic glutamate receptors induces apoptotic hair-cell death independent of afferent and efferent innervation. Sci Rep 7, 41102.

Sheets, L., Trapani, J.G., Mo, W., Obholzer, N., and Nicolson, T. (2011a). Ribeye is required for presynaptic Ca(V)1.3a channel localization and afferent innervation of sensory hair cells. Development 138, 1309–1319.

Sheets, L., Trapani, J.G., Mo, W., Obholzer, N., and Nicolson, T. (2011b). Ribeye is required for presynaptic Ca(V)1.3a channel localization and afferent innervation of sensory hair cells. Development (Cambridge, England) 138, 1309–19.

Sheets, L., Kindt, K.S., and Nicolson, T. (2012). Presynaptic CaV1.3 channels regulate synaptic ribbon size and are required for synaptic maintenance in sensory hair cells. J. Neurosci. 32, 17273–17286.

Sheets, L., Hagen, M.W., and Nicolson, T. (2014). Characterization of Ribeye Subunits in Zebrafish Hair Cells Reveals That Exogenous Ribeye B-Domain and CtBP1 Localize to the Basal Ends of Synaptic Ribbons. PLOS ONE 9, e107256.

Sheets, L., He, X.J., Olt, J., Schreck, M., Petralia, R.S., Wang, Y.-X., Zhang, Q., Beirl, A., Nicolson, T., Marcotti, W., et al. (2017). Enlargement of Ribbons in Zebrafish Hair Cells Increases Calcium Currents But Disrupts Afferent Spontaneous Activity and Timing of Stimulus Onset. J. Neurosci. 37, 6299–6313.

Sheng, Z.-H., and Cai, Q. (2012). Mitochondrial transport in neurons: impact on synaptic homeostasis and neurodegeneration. Nature Reviews Neuroscience 13, 77–93.

Sidi, S., Busch-Nentwich, E., Friedrich, R., Schoenberger, U., and Nicolson, T. (2004). gemini encodes a zebrafish L-type calcium channel that localizes at sensory hair cell ribbon synapses. J. Neurosci. 24, 4213–4223.

Song, H., Nie, L., Rodriguez-Contreras, A., Sheng, Z.-H., and Yamoah, E.N. (2003). Functional interaction of auxiliary subunits and synaptic proteins with Ca(v)1.3 may impart hair cell Ca2+ current properties. J. Neurophysiol. 89, 1143–1149.

Song, Q., Shen, P., Li, X., Shi, L., Liu, L., Wang, J., Yu, Z., Stephen, K., Aiken, S., Yin, S., et al. (2016). Coding deficits in hidden hearing loss induced by noise: the nature and impacts. Scientific Reports 6, 25200.

Srivastava, S. (2016). Emerging therapeutic roles for NAD+ metabolism in mitochondrial and age-related disorders. Clin Trans Med 5, 25.

Stankiewicz, T.R., Gray, J.J., Winter, A.N., and Linseman, D.A. (2014). C-terminal binding proteins: central players in development and disease. Biomolecular Concepts 5, 489–511.

Thevenaz, P., Ruttimann, U.E., and Unser, M. (1998). A pyramid approach to subpixel registration based on intensity. IEEE Transactions on Image Processing 7, 27–41.

Thio, S.S.C., Bonventre, J.V., and Hsu, S.I.-H. (2004). The CtBP2 co-repressor is regulated by NADH-dependent dimerization and possesses a novel N-terminal repression domain. Nucleic Acids Res 32, 1836–1847.

Todorova, V., and Blokland, A. (2017). Mitochondria and Synaptic Plasticity in the Mature and Aging Nervous System. Current Neuropharmacology 15, 166–173.

tom Dieck, S., Altrock, W.D., Kessels, M.M., Qualmann, B., Regus, H., Brauner, D., Fejtová, A., Bracko, O., Gundelfinger, E.D., and Brandstätter, J.H. (2005). Molecular dissection of the photoreceptor ribbon synapse. The Journal of Cell Biology 168, 825–836.

Tritsch, N.X., Yi, E., Gale, J.E., Glowatzki, E., and Bergles, D.E. (2007). The origin of spontaneous activity in the developing auditory system. Nature 450, 50–55.

Tritsch, N.X., Rodríguez-Contreras, A., Crins, T.T.H., Wang, H.C., Borst, J.G.G., and Bergles, D.E. (2010). Calcium action potentials in hair cells pattern auditory neuron activity before hearing onset. Nat. Neurosci. 13, 1050–1052.

Tucker, T., and Fettiplace, R. (1995). Confocal imaging of calcium microdomains and calcium extrusion in turtle hair cells. Neuron 15, 1323–1335.

Vos, M., Lauwers, E., and Verstreken, P. (2010). Synaptic mitochondria in synaptic transmission and organization of vesicle pools in health and disease. Front Synaptic Neurosci 2, 139.

Wang, X., Zhu, Y., Long, H., Pan, S., Xiong, H., Fang, Q., Hill, K., Lai, R., Yuan, H., and Sha, S.-H. (2018). Mitochondrial Calcium Transporters Mediate Sensitivity to Noise-Induced Losses of Hair Cells and Cochlear Synapses. Front Mol Neurosci 11, 469.

Xu, W., and Lipscombe, D. (2001). Neuronal Ca(V)1.3alpha(1) L-type channels activate at relatively hyperpolarized membrane potentials and are incompletely inhibited by dihydropyridines. J. Neurosci. 21, 5944–5951.

Yamoah, E.N., Lumpkin, E.A., Dumont, R.A., Smith, P.J., Hudspeth, A.J., and Gillespie, P.G. (1998). Plasma membrane Ca2+-ATPase extrudes Ca2+ from hair cell stereocilia. J. Neurosci. 18, 610–624.

Yin, Y., Liberman, L.D., Maison, S.F., and Liberman, M.C. (2014). Olivocochlear innervation maintains the normal modiolar-pillar and habenular-cuticular gradients in cochlear synaptic morphology. Journal of the Association for Research in Otolaryngology : JARO 15, 571–83.

Zenisek, D., and Matthews, G. (2000). The role of mitochondria in presynaptic calcium handling at a ribbon synapse. Neuron 25, 229–237.

Zhang, L., Engler, S., Koepcke, L., Steenken, F., and Köppl, C. (2018a). Concurrent gradients of ribbon volume and AMPA-receptor patch volume in cochlear afferent synapses on gerbil inner hair cells. Hearing Research 364, 81–89.

Zhang, Q., Li, S., Wong, H.-T.C., He, X.J., Beirl, A., Petralia, R.S., Wang, Y.-X., and Kindt, K.S. (2018b). Synaptically silent sensory hair cells in zebrafish are recruited after damage. Nat Commun 9, 1388.

